# Comprehensive enhancer-target gene assignments improve gene set level interpretation of genome-wide regulatory data

**DOI:** 10.1101/2020.10.22.351049

**Authors:** Tingting Qin, Christopher Lee, Raymond Cavalcante, Peter Orchard, Heming Yao, Hanrui Zhang, Shuze Wang, Snehal Patil, Alan P Boyle, Maureen A Sartor

## Abstract

Revealing the gene targets of distal regulatory elements is challenging yet critical for interpreting regulome data. Experiment-derived enhancer-gene links are restricted to a small set of enhancers and/or cell types, while the accuracy of genome-wide approaches remains elusive due to the lack of a systematic evaluation. We combined multiple spatial and *in silico* approaches for defining enhancer locations and linking them to their target genes aggregated across >500 cell types, generating 1,860 human genome-wide distal **En**hancer to **T**arget gene **Def**initions (**EnTDefs**). To evaluate performance, we used gene set enrichment testing on 87 independent ENCODE ChIP-seq datasets of 34 transcription factors (TFs) and assessed concordance of results with known TF Gene Ontology (GO) annotations., assuming that greater concordance with TF-GO annotation signifies better enrichment results and thus more accurate enhancer-to-gene assignments. Notably, the top ranked 741 (40%) EnTDefs significantly outperformed the common, naïve approach of linking distal regions to the nearest genes (FDR < 0.05), and the top 10 ranked EnTDefs performed well when applied to ChIP-seq data of other cell types. These general EnTDefs also showed comparable performance to EnTDefs generated using cell-type-specific data. Our findings illustrate the power of our approach to provide genome-wide interpretation regardless of cell type.

## Background

Enhancers, silencers and insulators are key genomic cis-regulatory elements that play pivotal roles in spatiotemporal control of gene expression by physical contact with the promoters of target genes they control [1–3]. Promoters are located immediately upstream of the transcription start sties (TSSs), facilitating the recruitment of transcription factors and RNA polymerase II (RNAPII) to instruct the initiation and direction of gene transcription, whereas enhancers and silencers can be located anywhere in the genome and often at distal regions, such as upstream, downstream or in introns of target genes or unrelated genes. Via interaction with promoters of the target genes, enhancers are bound by activator proteins and stimulate the rate of transcription, while silencers were bound by repressor proteins and decrease the rate. In certain cases when the interactions between enhancers/silencers and promoters are unwanted, insulators can block their interactions[4]. Bound by tissue-specific transcription factors and cofactors, such as p300 and Mediator, the cis-regulatory elements and promoter connections direct what, when and how the genome is transcribed so as to control cell fate decisions during development and differentiation [5–7]. For simplicity, we will refer to these distal cis-regulatory elements as general “enhancers” (>5kb from a transcription start site [TSS]) hereinafter.

Perturbation of enhancer activities and/or functions induced by genomic variants, epigenomic dysregulation, and/or aberrant chromosomal rearrangements can underlie disease susceptibility and developmental malformations [8, 9]. A prototypic example of this is the point mutation in the *Shh* enhancer, ZRS *(zone of polarizing activity regulatory sequence),* which can lead to limb malformations such as polydactyly in humans [10]. Recently, genome-wide association studies (GWAS) identified that >88% of disease-linked variants occur within non-coding regulatory DNA [11], especially enriched in enhancers [12]. These findings confirm the importance of enhancers in orchestrating transcriptional regulation and reveal that the dysregulation of enhancer function contributes to the pathogenesis of a variety of diseases, referred to as “enhanceropathies” [13].

A challenge in enhancer biology is to decipher their target genes and the mechanisms underlying the precise enhancer-gene interactions, which is reviewed in Pennacchio LA *et al* [14]. The enhancer to target gene specificity is essential to understand how gene expression is programmed during normal development and differentiation, and how the ectopic enhancer and/or non-target gene interactions can lead to diseases. However, interpreting genome-wide regulatory data is significantly hampered by our limited knowledge of enhancers and their target genes for multiple reasons. First, enhancers are commonly located distal to their target genes with multiple intervening genes in between, and greatly varying distances. One enhancer can act on multiple genes and one gene can be regulated by multiple enhancers [15]. Second, enhancers act in a dynamic and often cell type-specific manner, which further complicates the definition of a comprehensive set of enhancers and their target genes. Third, enhancers and promoters share various characteristics and functions [16, 17], thus making it challenging to disentangle the two elements based on functional genomic data.

With the breathtaking progress in technologies such as massive parallel sequencing and high-resolution chromosome conformation capture, our knowledge of cis-regulatory elements’ function and spatial organization have grown considerably over the past decade [18–23]. In most cases, enhancers are located at regions distal of their target genes up to hundreds of kilobases, and they can bypass more proximally located genes to bind to the promoters of the genes they control through long-range 3D chromosomal interactions [19, 24, 25]. The 3D genome is organized in hierarchical layers, from bottom to top including chromatin loops (or insulated neighborhoods), topological associating domains (TADs), and compartments [26]. The chromatin loops are the fundamental structural and functional building blocks of genome organization, which form between two convergent CTCF (CCCTC binding factor) binding sites bound by the cohesin protein complex [27].

Large epigenomics consortia like ENCODE [28–30] and Roadmap Epigenomics [31], have generated a tremendous amount of regulatory data across various tissue and cell types, including genome-wide transcription factor (TF) binding by ChIP-seq [32], chromatin accessibility assays (e.g. DNase-seq [33], ATAC-seq [34]), genome-wide chromatin mark profiles, and 3D chromosome organization. However, enhancer-promoter interactions are still highly restricted to a small number of cell types, which is probed by Chromatin Interaction Analysis by Paired-End Tag Sequencing (ChIA-PET [35]), and the genome-wide interaction map is still limited due to the high cost of Hi-C experiments [36]. Other enhancer-promoter interaction datasets have been generated by mathematical and/or bioinformatic approaches. The FANTOM5 [37] dataset is based on the gene expression correlation between enhancer and promoter regions, and Thurman et al. exploited DNase signal correlation between enhancers and promoters using DNase-seq data [38]. However, the reliability and generalization of these approaches remains elusive due to the lack of a systematic evaluation.

Gene set enrichment (GSE) testing is widely applied to infer the regulatory networks embedded in the abundant high-throughput gene regulation data, including ChIP-seq, Bisulfite sequencing, DNase-seq, and ATAC-seq. The first step in this analysis is to assign the genomic regions identified by the assays to their target genes, and most methods simply do the assignment using the nearest genes regardless of the actual regulatory targets [39–42]. Since enhancers and their target genes have long-range chromosomal contact, adjacent gene assignments tend to link enhancers to non-target genes, leading to incorrect interpretation for distal enhancer regulation. In this study, we aimed to determine the best sets of human “enhancers” (enhancers, silencers and insulators) and their gene targets. By all possible combinations of existing experimental and/or computationally-derived datasets, we generated 1,860 **En**hancer to **T**arget gene **Def**initions, referred to as EnTDefs, and systematically evaluated their performance based on the concordance of GSE results of 87 ENCODE ChIP-seq datasets with known TF biological processes, resulting in a handful of best-performing EnTDefs. We also showed that as opposed to being random, target genes that are often missed or often falsely identified using adjacent gene assignments are biased to specific Gene Ontology terms. In addition, we compared cell-type-specific EnTDefs (CT-EnTDefs) with non-cell-type-specific ones (general EnTDefs) and found that general EnTDefs were more favorable. Our findings demonstrate that the novel, top-performing EnTDefs significantly enhance the biological interpretation for genomic region data regardless of cell type.

## RESULTS

### Creation and ranking of genome-wide Enhancer-to-Target gene Definitions (EnTDefs)

Several approaches to define human enhancer locations and their target genes have been proposed in the literature, but no systematic study has been performed to evaluate their performance separately or in combination on a genome-wide scale. To determine the best sets of human enhancers and their distal gene targets, we generated a total of 1,860 genome-wide Enhancer-Target gene Definitions (EnTDefs) using existing experiments and/or literature-derived data, and systematically evaluated their performance. This was done by applying all possible combinations of methods for defining *1) enhancer region locations,* identified from four data sources (ChromHMM[43], DNase-seq[38], FANTOM5[37, 44, 45] and Thurman[38]), and *2) enhancer-target gene links,* defined by four different methods (ChIA-pet data [“ChIA”][46, 47], DNAase-signal correlation [“Thurman”][38], gene expression correlation [“FANTOM5”][45] and loop boundaries with convergent CTCF motif [“L”][48]), including combinations using multiple of each (see Methods for details). Overall, these included a total of 1,768,201 possible individual enhancer-target links across >500 cell types by integrating all of the 4 enhancer-defining datasets and all of the 4 enhancer-gene link datasets. These enhancer-target links were defined from 685,921 enhancers and 21,094 linked target genes. **Figure 1** demonstrates the workflow for the creation and evaluation of these 1,860 EnTDefs. For the “L” enhancer-gene linking method, we evaluated the loops with up to 3 genes (L1: one gene, L2: ≤ two genes, or L3: ≤ three genes), allowing the links between the enhancer to any of the included genes within the loop. Because current knowledge of enhancers is far from complete and the experimental data that assay enhancers to target genes is limited, the genome coverage of EnTDefs defined by the experimentally and/or computationally derived methods (**Figure 1A:** four enhancer-defining methods and four enhancer-target gene linking methods) was expected to be low. Therefore, we extended the enhancer regions up to 1kb and/or assigned regions outside of enhancers and promoters (within 5kb of a transcription start site (TSS)) to the gene with the nearest TSS (**Figure 1A:** Extension and Additional link), resulting in 100% coverage of distal genomic regions (>5kb of TSS). All of the 1,860 EnTDefs were evaluated and ranked based on how well they performed in gene set enrichment (GSE) testing with genes’ distal ChIP-seq peaks. Specifically, the Gene Ontology biological process (GO BP) enrichment results from 87 ENCODE ChIP-seq datasets for 34 distinct transcription factors (TFs) were compared with the curated GO BP terms annotated to the same tested TFs (GO annotation by GO database) using F1 scores (*see Methods).* EnTDefs demonstrating higher concordance ranked higher, as they were better able to identify the known functions of the TFs based on their distal binding regions (non-promoters).

**Figure 1.**
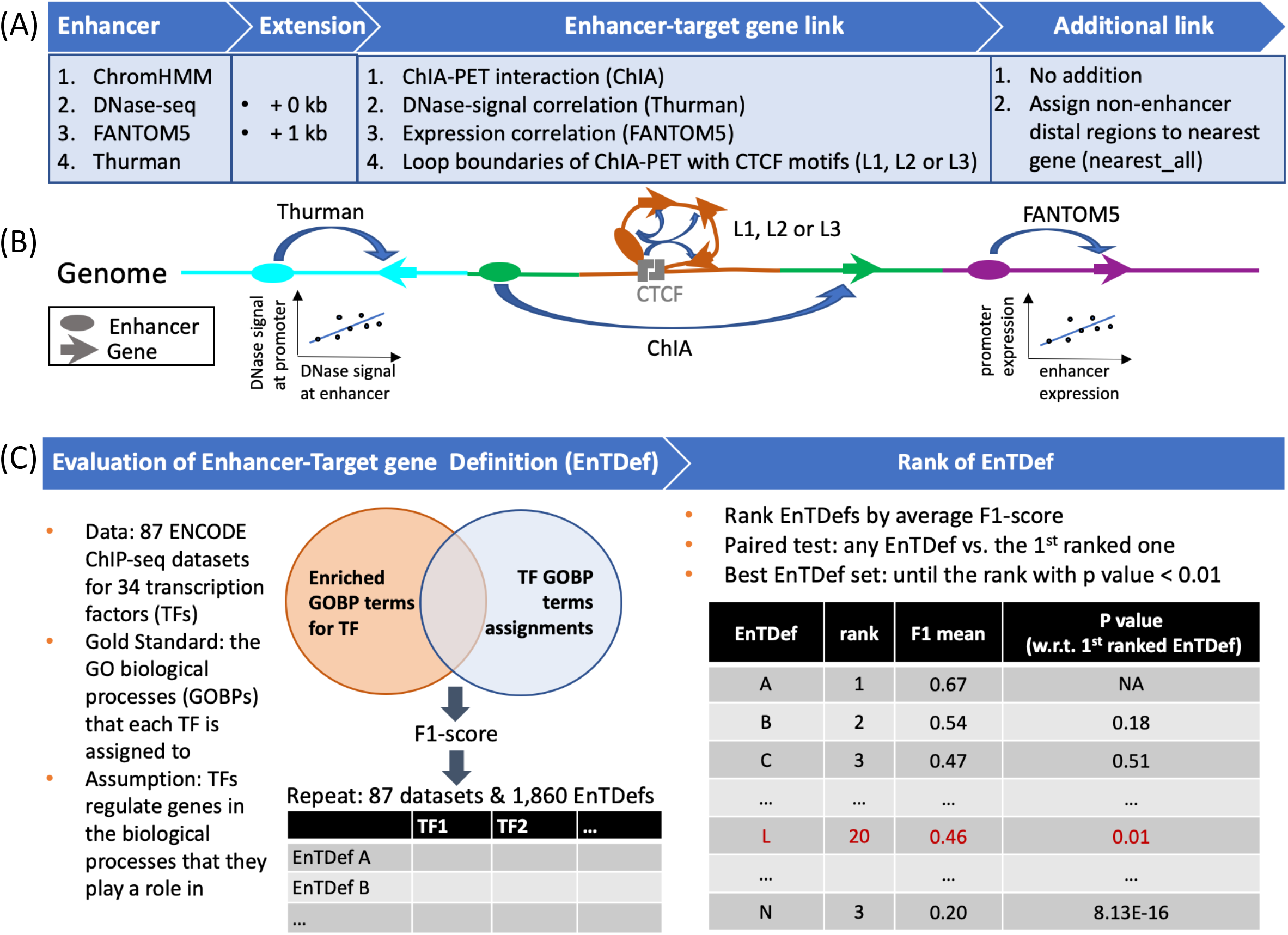
Workflow for generating and evaluating EnTDefs. (A) Enhancers were defined by ENCODE ChromHMM UCSC tracks, ENCODE DNase-seq hypersensitive sites (DHSs), Cap Analysis Gene Expression (CAGE) experiment-derived enhancers from the FANTOM5 project, and/or distal and non-promoter DHS within 500kb of the correlated promoter DHSs from Thurman *et al.* (B) The enhancer-target gene links were defined by ChIA-PET interactions from ENCODE ChIA-PET data (ChIA), DNase-signal correlation-based links from Thurman *et al,* expression correlationbased interactions from FANTOM5, and/or interactions between enhancers and genes within loop (L) boundaries of ChIA-PET with convergent CTCF motifs (L1 [one gene], L2 [≤ two gene] or L3 [≤ three genes] were allowed). An enhancer can be assigned to multiple genes. To increase the genome coverage, we allowed the extension of enhancers to 1kb (i.e. enhancer extension), and assigned other regions outside of 5kb from a TSS to the nearest gene (i.e. “nearest_All” additional links). All combinations of the above, allowing multiple at a time, defined the possible **En**hancer to **T**arget gene **Def**initions (EnTDefs). (C) Left: 1,860 EnTDefs were generated and GOBP GSE testing was performed on 87 ENCODE TF ChIP-seq datasets using each of the EnTDefs. By comparing the significant GOBP terms identified by GSE with each EntDef to those assigned to the TF by the GO database (“GO annotation”), the F1-score was calculated for each EnTDef-TF pair. Right: the EnTDefs were ranked by average F1-score across TFs in descending order. TF paired Wilcoxon sum-rank test was performed between the top ranked EnTDef and each of the sequential ones to identify the set of best EnTDefs (top 1 until the rank with p value < 0.01).

### Overview of the EnTDef characteristics

We first investigated the characteristics of the 1,860 EnTDefs by comparing them to simply assigning distal genomic regions (i.e. >5kb from a TSS) to the genes with the nearest TSS (>5kb **Loc**us **Def**inition [LocDef]) (**Figure 2A, Supplementary Figure S1**). The EnTDefs were ranked in decreasing order by their average F1 score across 34 TFs, and the top 741 EnTDefs (~40%) were found to significantly outperform the >5kb LocDef (Wilcox signed-rank test, FDR < 0.05). The best performing EnTDef (No. 1 ranked) was defined by DNase-seq plus FANTOM5 enhancers and ChIA, Thurman and FANTOM5 enhancer-target gene link methods with the “nearest_All” addition. For the top 741 EnTDefs, the percentage of genome covered and percent of distal peaks caught (outside of 5kb regions around TSSs) was as high as 100% (89% – 100%), the median number of genes assigned to each enhancer was 2 (range of 1 – 2), and the median number of enhancers assigned to each gene was 20 (range of 2 – 98). Out of the 741 EnTDefs, those ranked 2 through 19 were not significantly worse than the best performing EnTDef (Wilcoxon signed-rank test, p > 0.01. **Supplementary Table S1**), suggesting that these 19 EnTDefs performed equally well. This finding was robust to the specific set of GOBP annotations used (i.e. with or without IEA-based GO to gene annotations; see *Methods,* data not shown).

**Figure 2.**
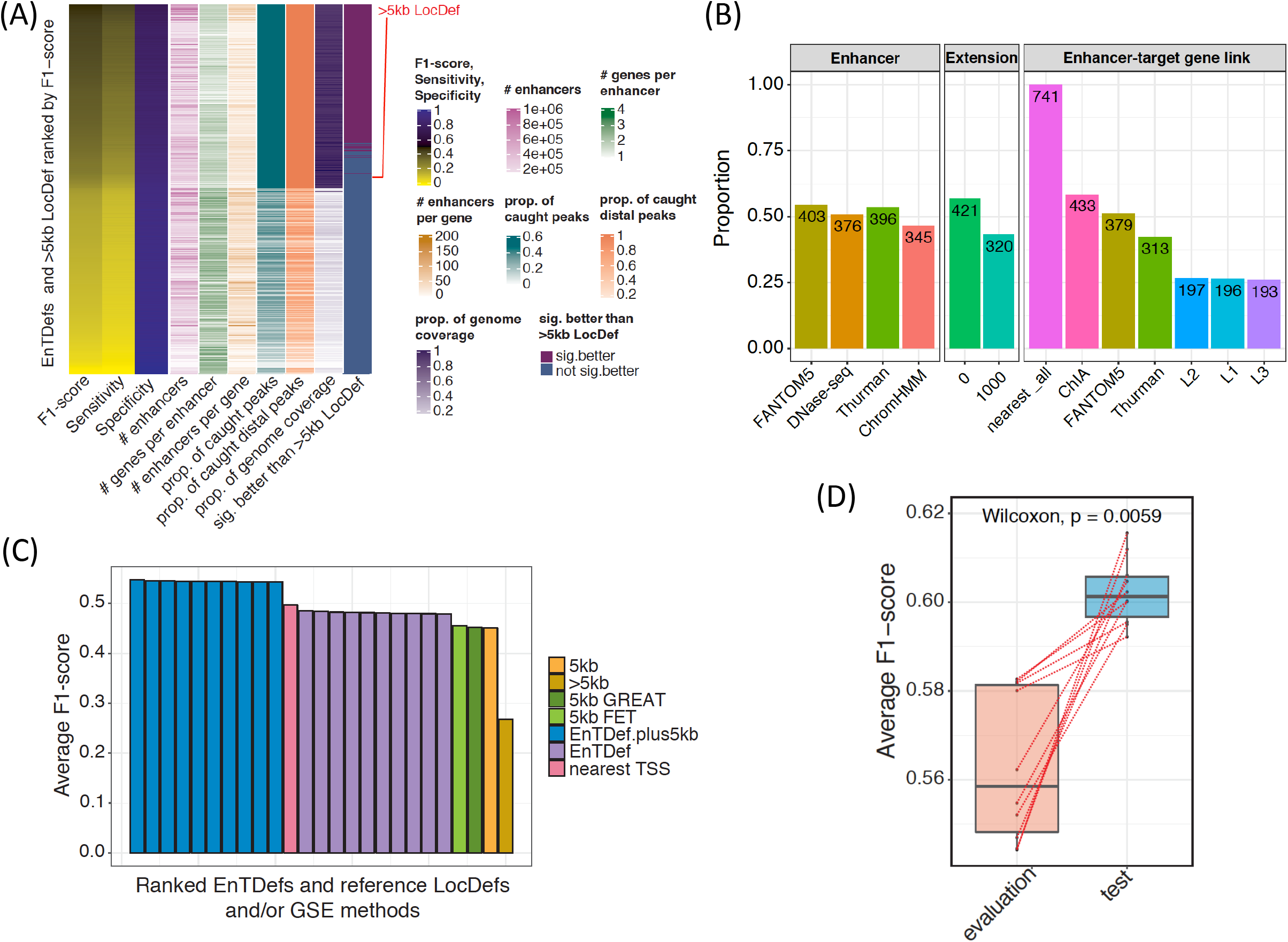
Characteristics of EnTDefs. (A) Overview of the characteristics of 1,860 EnTDefs and >5kb LocDef ranked by F1-score in descending order. F1-score, sensitivity, specificity, number of enhancers, average number of genes per enhancer, average number of enhancers per gene, average proportion of caught TF peaks, average proportion of caught TF peaks outside of 5kb of TSSs, proportion of genome coverage, and whether the EnTDef was significantly better than the >5kb LocDef are shown. (B) The frequency of each method (four enhancer definition methods, with or without enhancer extension, seven enhancer-target gene link methods) among the 741 EnTDefs that significantly outperformed the >5kb LocDef. For simplification, the “nearest_all” additional link method was grouped in enhancer-target gene link method. (C) Bar plot of average F1-scores for the top 10 EnTDefs plus 5kb LocDef (blue bars), top 10 EnTDefs (purple bars), nearest TSS method (pink bar), >5kb LocDef (mustard bar) and 5kb LocDef used by PE.Approx (yellow bar), by GREAT (dark green bar) and by Fisher’s exact test (green bar). (D) Distribution of average F1-scores of the top 10 EnTDefs used on *evaluation* ChIP-seq datasets or *testing* ChIP-seq datasets. The dashed lines link the same EnTDefs used in the two different ChIP-seq datasets. The p value of Wilcoxon signed-rank test was shown in the figure.

By examining the types of methods used to generate the top 741 EnTDefs (**Figure 2B**), we found that: *i)* over half of them included FANTOM5 (54%), Thurman (53%) and DNase-seq (51%) enhancer regions, while chromHMM defined enhancers were least used (~45%); *ii)* approximately 60% of them were generated without enhancer regions extension; *iii)* all of them used the “nearest_All” addition to assign the distal regions that were not in enhancers to the nearest gene’s TSS; and *iv)* the “ChIA” method applying ChIA-PET data to assign enhancers to target genes was included most frequently (~60%), followed by FANTOM5 (~51%) and Thurman (42%), whereas the “L” method assigning genes to enhancers within the same loop boundaries with convergent CTCF motifs was least used (~26%). It is not surprising that all of the top 741 EnTDefs included the “nearest_all” addition, because this significantly increased genome coverage by assigning all regions outside enhancers and promoters to the nearest distance gene (>5kb LocDef), leading to improved sensitivity and thus F1 score (**Figure 2A**). On the other hand, the fact that these 741 EnTDefs outperformed the >5kb LocDef suggests that the “smart” enhancer to target gene assignments more accurately capture real biological regulatory elements for distal enhancer regions when compared to the simplistic assignment to nearest genes. In addition, all of the top performing EnTDefs were generated using combinations of at least two different datasets/methods for enhancer definitions (ChromHMM, DNase-seq, FANTOM5, and/or Thurman) and enhancer-gene assignments (ChIA, FANTOM5, L, and/or Thurman), illustrating the importance of high genomic coverage and that the integration of multiple data sources and methods indeed improves the performance of enhancer to target gene assignments.

### EnTDefs plus promoter regions outperforms the nearest TSS method

Our analyses thus far have focused on the assessment of distal gene regulation. However, often the goal is to assess the functional regulation from anywhere in the genome, including binding both distal and proximal to TSSs. One commonly used method for ChIP-seq GSE testing is to link all peaks to the nearest gene, hereinafter referred to as the “nearest TSS” method (**Supplementary Figure S1**: “nearest TSS” LocDef), resulting in all peaks having at least one assigned gene. EnTDefs were generated for distal regions (outside the 5kb windows around TSSs) and any regions within 5kb of a TSS were ignored, whereas the “nearest TSS” method includes all genomic regions. Thus, to compare fairly with the “nearest TSS” method, we added promoter regions to the top 10 ranked EnTDefs, referred to as “EnTDef_plus5kb”. That is, peaks within 5kb of a TSS were assigned to the nearest gene (**Supplementary Figure S1**: “5kb” LocDef), while distal peaks were assigned according to the EnTDef. All ten of the EnTDef_plus5kbs significantly outperformed the “nearest TSS” method (~0.05 increase in average F1 score, Wilcoxon signed-rank test, p < 0.0001) (**Figure 2C**), using the same evaluation method based on F1 scores as used above (see *Methods).*

We next determined if our ‘smart’ EntDefs using only distal binding events could even outperform the use of all peaks (promoter and enhancer) with naïve assignments to the genes with the nearest TSS. When compared with the “nearest TSS” method, the top 10 best performing EnTDefs showed slightly lower F1 scores (~0.03 lower), but the difference among the top half of them were not significantly different from “nearest TSS” (Wilcoxon signed-rank test, p > 0.05). Thus, although they did not outperform it, the best were not significantly worse. This illustrates the great importance of regulation from promoters in GSE testing.

Two other commonly used GSE methods for genomic regions, GREAT[39] and Fisher’s exact test (FET) using peaks within 5kb of a TSS (**Supplementary Figure S1**: “5kb” LocDef), were also evaluated using the same scheme. Notably, the three GSE testing methods (Poly-Enrich, GREAT and FET using 5kb LocDef to assign peak to gene) performed equally well (Friedman test, p = 0.91), but significantly worse than the top 10 EnTDefs (Figure 2C, average F1 = 0.45 vs 0.47, Wilcoxon rank-sum test, p < 0.007). In addition, both top 10 EnTDefs and 5kb LocDef (i.e. assigning promoters to the nearest gene) significantly outperformed >5kb LocDef (i.e. the naïve approach of assigning distal regions to the nearest gene) (average F1 = 0.47, 0.45 vs 0.27, Wilcoxon signed-rank test, p = 2.37×10^-14^ and 1.32×10^-8^ respectively). In summary, although the naïve approach of linking distal regions to the nearest gene (>5kb LocDef) did not outperform the use of promoter data only (5kb LocDef), the use of distal binding events with ‘smart’ gene assignments (EnTDefs) did outperform the use of promoter data only. These findings illustrate the importance of accurately modeling regulation from enhancers, and that when done well, enhancers have the potential to provide more regulatory information than promoters. We conclude that GSE testing using our top EnTDefs exceeds the commonly used nearest distance-based and promoter-only based GSE approaches.

### Our EnTDefs are generalizable to different cell lines

Next, we sought to investigate whether the EnTDefs (which were selected based on their performance in GM12878, H1-HESC and K562 cell lines) can perform equally well on testing ChIP-seq data from different cell lines (A549, HEPG2, HUVEC and NB4). Surprisingly, the average F1 score of the top 10 EnTDefs across the *test* ChIP-seq datasets (different cell lines) was significantly higher than that from the *evaluation* ChIP-seq datasets (average F1 = 0.60 vs. 0.56, Wilcoxon sum-rank test, p = 0.0059) (**Figure 2D, and Supplementary Figure S2**). This may be due to the test ChIP-seq datasets containing more peaks than the evaluation datasets (**Supplementary Figure S3A**, Wilcoxon sum-rank test, p = 0.092), and indeed we found that the *F1 scores* were significantly corelated with the number of peaks (**Supplementary Figure S3B**, Pearson’s correlation r = 0.65, p = 4.57×10^-6^). The findings indicate that the performance of the top selected EnTDefs are independent of the cell types of ChIP-seq datasets, but likely strongly influenced by the quality of the datasets themselves. We reasoned that the EnTDefs were created based on the combinations of diverse data sources stemming from >500 different cell types, resulting in a consensus set of enhancer and gene assignments across various cell types, and therefore representative of the background interactions between enhancer and target genes across many cell types. The high generalizability of our top EnTDef makes it feasible to integrate with GSE testing in a cell type-independent manner.

### General EnTDefs perform comparably to cell-type-specific EnTDefs

To contrast with the EnTDefs generated by integrating data for many cell types, hereafter called “general EnTDefs”, we created “cell-type-specific EnTDefs” (CT-EnTDef) using ChIA-PET datasets of a particular cell type. Since many enhancers and regulatory relationships between enhancer and target genes are considered to be tissue and cell-type-specific, we sought to examine how the general EnTDefs perform when compared with CT-EnTDefs. For each tested TF (the average number of TFs tested in each cell type is ~60 [6–104], see **Supplementary Table S2**), F1 scores were calculated (see *Methods)* and compared between a pair of EnTDefs which were created using the same methods (the same combinations of enhancer and enhancer-gene link methods) but based on data from different cell types: *i)* general EnTDef vs. CT-EnTDef using the same cell type (same-CT-EnTDefs), *ii)* general EnTDef vs. CT-EnTDef using a different cell type (diff-CT-EnTDefs), and *iii)* same CT-EnTDefs vs. different CT-EnTDefs. Notably, there was no significant difference in F1 scores among the three comparative EnTDefs for cell types GM12878, H1HESC or K562 (**Figure 3A**, Kruskal-Wallis test, p ≥ 0.5). For MCF7, the same-CT-EnTDef performed near significantly better than the general CT-EnTDef (Wilcoxon sum-rank test, p = 0.03; three groups: Kruskal-Wallis test, p = 0.059). It is worth noting that only four TFs were tested in cell type MCF7, whereas ≥46 TFs were tested in the other three cell types, so this exception might not be visible for a larger set of TFs.

**Figure 3.**
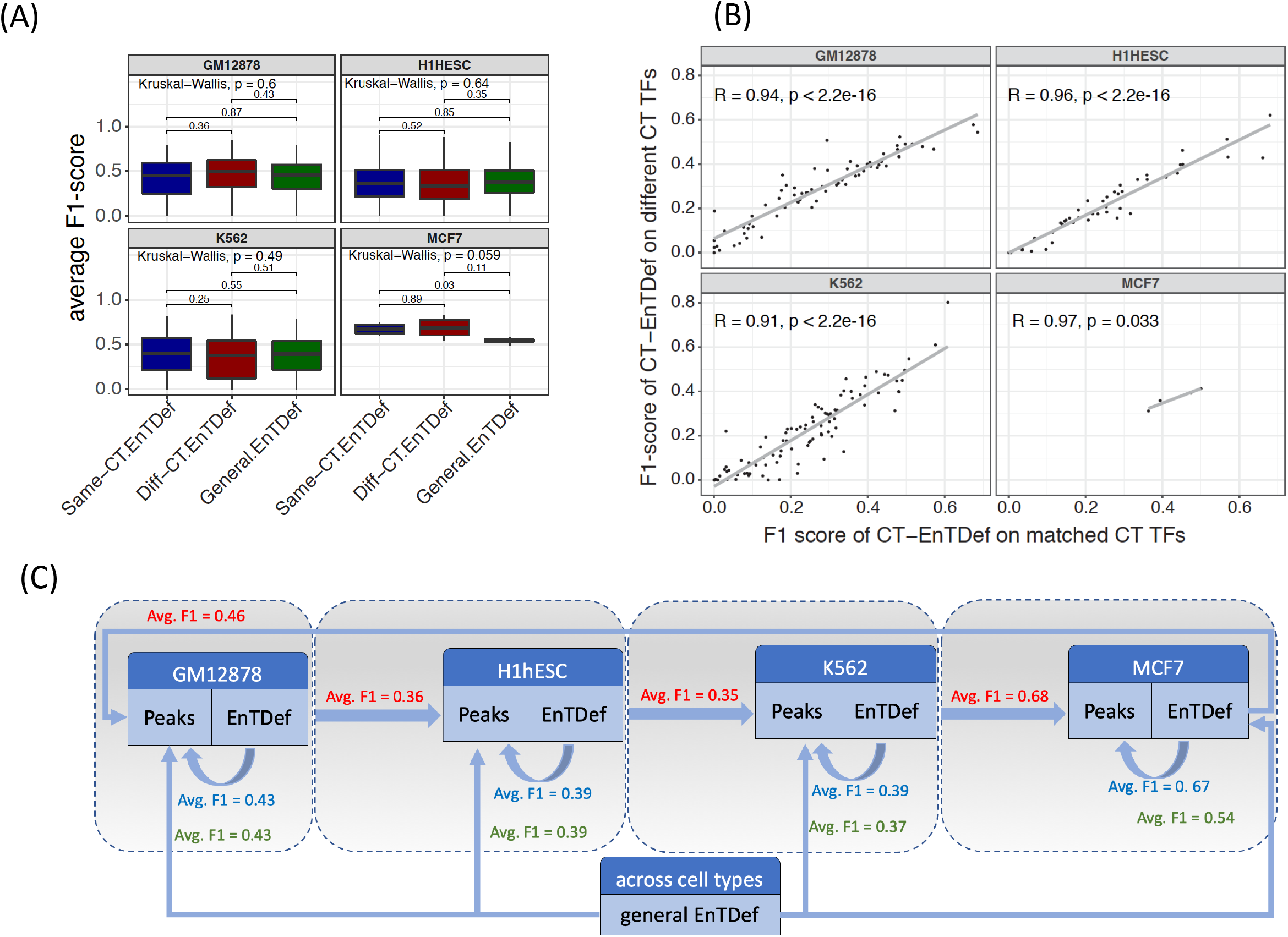
Evaluation of cell type-specific (CT)-EnTDefs and general EnTDefs. (A) Distribution of the average F1-scores of same-CT EnTDefs, diff-CT EnTDefs and general EnTDefs that were applied on the same TF ChIP-seq data. (B) Correlation between average F1-scores calculated on a TF in a particular cell type using CT-EnTDefs of the matched cell type (x-axis) and the ones calculated on the same TF using CT-EnTDefs of a different cell type (y-axis). Each dot represents an average F1-score of a TF across EnTDefs, and each panel is one of four cell types (GM12878, H1HESC, K562 and MCF7) for which the CT-EnTDefs were created and evaluated, respectively. (C) Evaluation summary of different types of EnTDefs in four different cell types. Comparative average F1-scores associated with a particular cell type are grouped in a grey box: blue refers to using the CT-EnTDef on a TF ChIP-seq from the same cell line, green refers to using the general CT-EnTDef on that TF ChIP-seq, and red refers to using the diff-CT-EnTDef on TF ChIP-seqs from that TF ChIP-seq.

We also observed that the average F1 scores of TFs across all possible EnTDefs were significantly correlated between the same-CT-EnTDef and diff-CT-EnTDef for all four cell lines (**Figure 3B and Supplementary Figure S4**, Pearson’s correlation > 0.9, p < 0.0001). This implies that the performance of the results is driven more by the quality and quantity of input data than by whether a general or CT-EnTDef is used. The trend of correlation still held at the individual TF and EnTDef level (F1 score per TF per EnTDef, **Supplementary Figure S5**). As shown in **Figure 3C**, regardless of the type of EnTDef (general EnTDefs, same-CT-EnTDefs and diff-CT-EnTDefs) used for evaluation, the average F1 score across all TFs and EnTDefs were similar, with the difference ranging from 0 to 0.14. Taken together, these findings suggest that CT-EnTDefs are overall comparable to general EnTDefs, and the benefit of using CT-EnTDefs is minor and depends on the quality and quantity of data for a particular cell type (e.g. MCF7 in **Figure 3C**). This is good news since it is costly and difficult to generate cell-type-specific ChIA-PET experiments, which are required to create the corresponding CT-EnTDef. In contrast, the general EnTDefs, which capture real enhancer and target gene interactions in a similar way to CT-EnTDef, are more practically and economically favorable for GSE testing.

### Incorrect gene assignments by nearest distance method are not random

Since enhancers are known to be located up to 1Mbp away from their regulatory genes [14, 49], several interceding genes can reside between a TF binding site (peak) in an enhancer and its target gene(s), as modeled by our EnTDefs (**Supplementary Figure S6**). In contrast, the nearest distance method simply links a peak to the gene with the nearest TSS without accounting for interceding genes. By ranking the genes based on the average number of interceding genes across the enhancers that target them, we investigated whether the number of interceding genes is randomly distributed across genes and GO terms, or if there are GO terms significantly enriched with genes having more or fewer interceding genes[50]. We investigated the best performing EnTDef excluding the “nearest_all” addition, in order to assess the ‘smart’ enhancer-target links only. The genes least likely to have interceding genes were found to be significantly enriched in *G protein-coupled receptor activity* (FDR = 1.41×10^-14^), *olfactory receptor activity* (FDR = 6.21×10^-12^), detection of chemical stimulus (FDR = 3.23×10^-11^)*, phenol-containing compound metabolic process* (FDR = 1.91×10^-4^)*, GABA-ergic synapse* (FDR = 2.35×10^-4^), *RISC complex* (FDR = 2.39×10^-4^), *postsynaptic membrane* (FDR = 4.13×10^-4^) and *behavior* (FDR =4.71×10^-4^) (**Figure 4A**). These GO terms enriched with genes least likely to have interceding genes (lower ranked genes) are most likely to be correctly assigned by the nearest distance method (**Supplementary Figure S1**: >5kb LocDef), and thus most easily detectable by current GSE testing. Conversely, the GO terms enriched with higher numbers of interceding genes (upper ranked genes) were *mRNA metabolic process* (FDR = 8.09×10^-8^), *regulation of catabolic process* (FDR = 8.40×10^-8^), *chromatin organization* (FDR = 2.53×10^-7^), *kinase binding* (FDR = 1.75×10^-6^), *heterocycle catabolic process* (FDR = 3.22×10^-6^), *chromatin* (FDR = 7.25×10^-6^), *hemopoiesis* (FDR = 9.47×10^-6^) and *RNA processing* (FDR = 2.27×10^-5^) (Figure 4A). Those GO terms are least likely to be assigned by the nearest distance method, and most likely missed using current methods for GSE testing.

**Figure 4.**
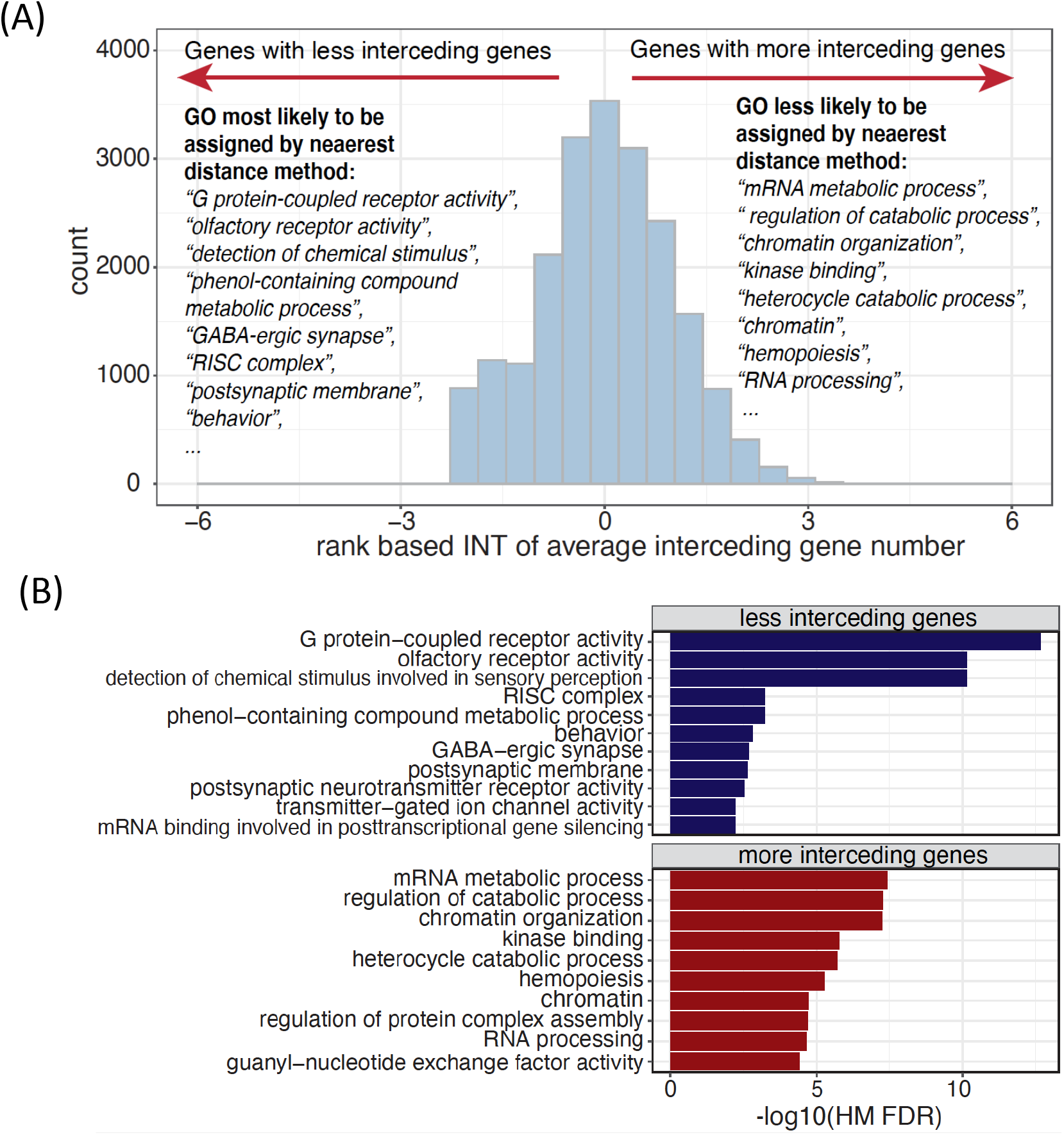
GO terms often missed or falsely identified by the nearest distance method of assigning genomic regions to target genes. (A) Distribution of the rank-based inverse normal transformation (INT) of average interceding gene numbers for the best EnTDef without the “nearest_all” addition. The top ranked enriched GO terms most likely or less likely to be identified by nearest distance method were listed. (B) The enriched GO terms in the genes with fewest interceding genes and the ones with the most interceding genes across the top 10 EnTDefs and their associated -log10 Harmonic Mean (HM) FDR.

To determine if this observation is robust to different EnTDefs, we performed the same analysis on all top 10 best performing EnTDefs without the “nearest_all” addition, and combined the results by calculating FDR-adjusted harmonic mean p-values, followed by removing redundant terms (see *Methods).* Consistently, *G protein-coupled receptor activity, olfactory receptor activity, RISC complex* and *postsynaptic membrane* were still the top 5 enriched terms for the genes with fewer interceding genes, and similarly, *regulation of catabolic process, chromatin organization, kinase binding* and *heterocycle catabolic process* were the top 5 enriched terms in upper ranked genes with more interceding genes (**Figure 4B**). These findings indicate that both the genes with the most and fewest interceding genes are not random: chemical stimulus and neuron-related genes can be easily assigned with the nearest distance method, whereas metabolic processing and chromatin organization genes may be frequently missed. It is concordant with the knowledge that enhancers regulate genes via long-range chromatin interactions, which are able to be captured by our EnTDefs.

### Nearest distance assignment method leads to false positive and false negative GSE results

Our finding that the nearest distance method tends to incorrectly assign TF binding sites in some biological functions more than others triggered us to investigate how this bias affects the results of GSE testing. By comparing the GSE results from assigning distal regions to the nearest genes (i.e. using >5kb LocDef) to those using the best-performing EnTDef, and evaluating using TF GO annotations as above (see Methods), we identified all false positive (FP) and false negative (FN) enriched GO terms by the >5kb LocDef, that were correctly called by our EnTDef. First, we ranked the FP GO terms in descending order by significance across the 34 tested TFs. These FPs represent GO terms with genes that tend to be in between enhancers and their targets (i.e. the interceding genes), and they are not annotated to the TFs of interest. For instance, the GO term *blood vessel morphogenesis* is annotated with *PROK1,* an interceding gene between an enhancer bound by the TF *YY1* (peak:1503) and its target gene *SLC16A4* (**Figure 5A left panel**). Using the nearest distance method, the *YY1* binding site would be falsely assigned to the nearest gene *PROK1*, rather than to the real target gene *SLC16A4,* leading to the enrichment of *blood vessel morphogenesis* that is not regulated by the TF *YY1* (i.e. a FP term by >5kb LocDef).

**Figure 5.**
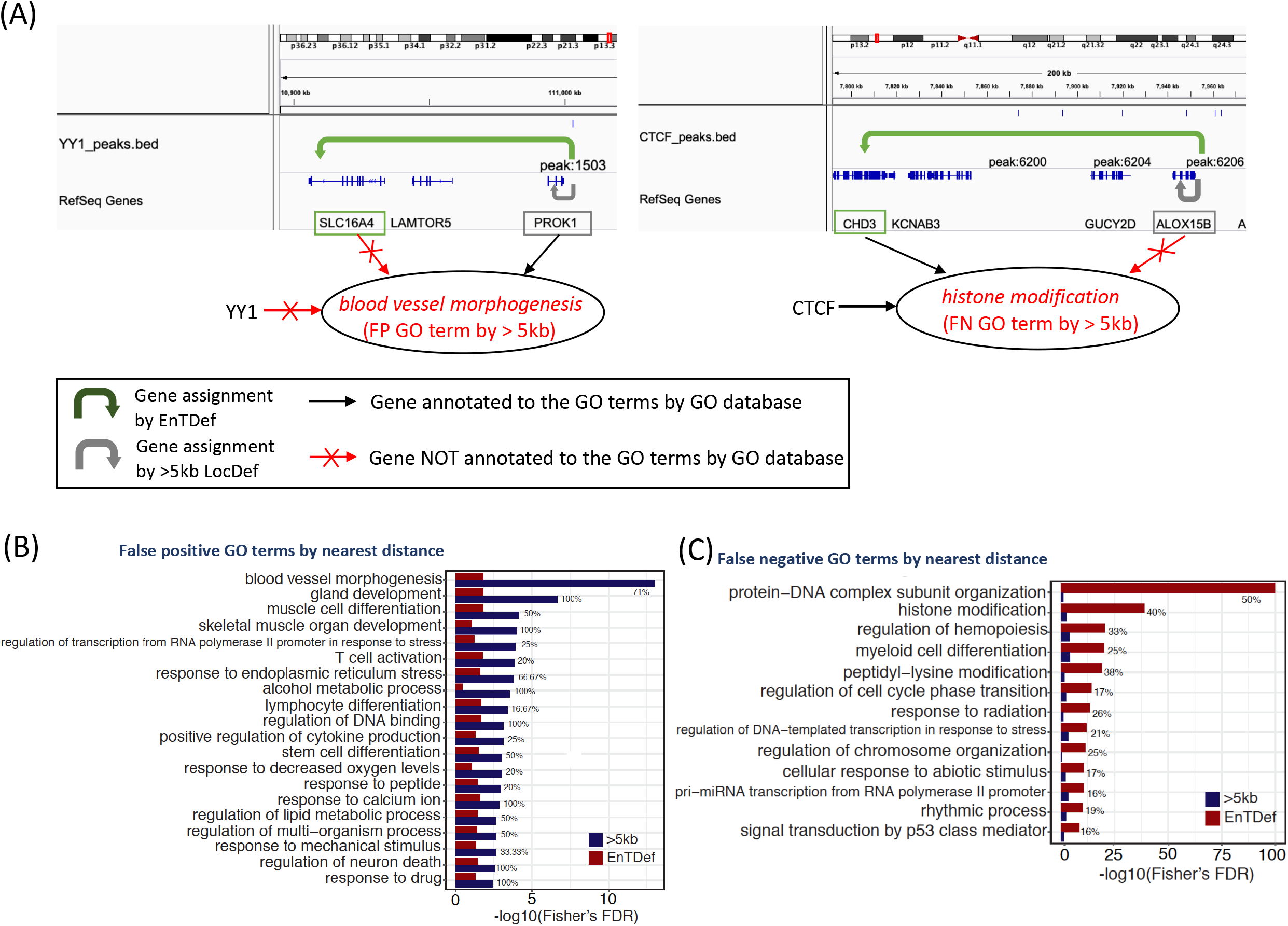
False positive and false negative GSE results by nearest distance assignment method. (A) Examples of false positive (left) and false negative results. The false positive term *blood vessel morphogenesis* was annotated with gene *PROK1,* the interceding genes between an enhancer bound by a TF *YY1* (peak:1503) and its target gene *SLC16A4.* The false negative term *histone modification* was annotated with gene *CHD3,* a target gene of the enhancer bound by the TF *CTCF* (peak:6206). (B) The false positive GO terms identified as enriched by the nearest distance method, but not by the best performing EnTDef and not defined by the TFs’ GO annotation. (C) The false negative GO terms which were failed to be identified by the nearest distance method, but correctly identified by the best performing EnTDef and also defined by the TFs’ GO annotation. The significance levels (-log10 Fisher’s FDR) are shown in dark blue for >5kb LocDef and dark red for the EnTDef. The numbers listed at the end of each paired bars are false positive rate (B) and false negative rate (C).

The top ranked FP biological processes (**Figure 5B**) included those related to development and regulation *(gland development, skeletal muscle organ development, regulation of DNA biding, and regulation of neuron death),* metabolic process or response to different stimuli *(alcohol metabolic process, response to calcium ion, response to drug),* and cell differentiation processes *(Stem cell differentiation, lymphocyte differentiation* and *T cell activation).* Among them, *blood vessel morphogenesis* was identified to be the most often falsely enriched term by >5kb LocDef (Fisher’s FDR = 9.48×10^-14^) with five out of seven TFs falsely called as positives (FP rate = 71%). It is noteworthy that the Fisher’s p values of FP terms calculated based on >5kb LocDef GSE results were overall significantly lower than those based on EnTDef GSE results (Wilcoxon rank-sum test, p= 3.95×10^-09^), indicating that those FP terms were much less likely identified by our EnTDef method.

On the other hand, common FN GO terms failed to be identified by >5kb LocDef while being successfully identified by the top EnTDef. In contrast to the FP GO terms, FN terms tend to be missed by assignment to the nearest distance gene, but correctly identified using the top EnTDefs. As shown in **Figure 5A** (right panel), the GO term *histone modification* contains the gene *CHD3*, the gene target of an enhancer bound by the TF *CTCF* (the peak:6206). Using the nearest distance method, the *CTCF* binding site would be falsely assigned to the nearest gene *ALOX15B,* rather than the target gene *CHD3,* failing to identify *histone modification* as regulated by the TF *CTCF* (i.e. a FN term by >5kb LocDef). These FN GO terms consistently point to chromosome organization and modification processes, including *Protein-DNA complex subunit organization* (FN rate = 50%), *histone modification* (FN rate = 40%), *Peptidyl-lysine modification* (FN rate = 38%), and *regulation of chromosome organization* (FN rate = 25%) (**Figure 5C**). Their Fisher’s p values based on the >5kb LocDef GSE results were significantly higher than those based on the EnTDef GSE results (Wilcoxon rank-sum test, p= 6.24×10^-15^), in line with the finding that chromosome organization-related terms are very often missed by the nearest distance method. Together, these results demonstrate that GSE analysis using the nearest distance gene assignment method cannot always identify biological processes induced by long-range chromosome organization; rather, they tend to favor development and cell differentiation functions which are not related to the TFs. In contrast, the EnTDef method can successfully detect distal enhancer and target gene interactions even for biological processes with complex long-range interactions such as chromosomal organization-related terms, and avert potential false positive results.

### Guidance for selecting a peak-to-gene assignment method in GSE analysis

The first step in GSE testing of cis-regulome data, such as TF binding sites or chromatin marks from ChIP-seq, is to assign the genomic regions or peaks to their target genes. The different assignment methods can lead to variable enrichment results and FP and/or FN findings, as discussed above (nearest distance method vs. EnTDef). To avoid misinterpretation of genomewide regulatory data, we need to select an appropriate LocDef method with care, which should be specific to the particular research question and the genomic regions of interest. **Figure 6** summarizes three general categories of research questions and the corresponding regions of interest: *i)* the 5kb or 1kb LocDef should be selected when interested in how a TF and/or chromatin mark regulates gene expression from promoters; *ii)* the EnTDef (enhancer) should be selected when interested in how a TF and/or chromatin mark regulates gene expression from distal regions; and *iii)* when the comprehensive regulatory signature is of interest, including both promoter and distal regions, our EnTDef plus 5kb LocDef (enhancer.5kb) should be selected. The promoter LocDef has the lowest genome coverage (10% for <5kb LocDef and 2% for <1kb LocDef), while the EnTDef plus 5kb has 100% genome coverage, and the EnTDef has intermediate genome coverage (90%). We incorporated our top performing EnTDef and EnTDef.plus5kb into the Bioconductor package *chipenrich* [42] and the ChIP-Enrich website (https://chip-enrich.med.umich.edu), allowing users to select the most suitable genomic regions-gene assignment methods, gene sets and GSE method to correctly interpret their genome-wide regulatory data. In addition, we provide a peak-to-gene assignment functionality in our GSE Suite (http://gsesuite.dcmb.med.umich.edu), by which users can select any possible combination of enhancer location and enhancer-to-gene target methods (as described in this study) and obtain the gene assignments for a user uploaded list of genomic regions, based on the selected EnTDef, or other method (e.g. promoters, exons, introns or anywhere in the genome).

**Figure 6.**
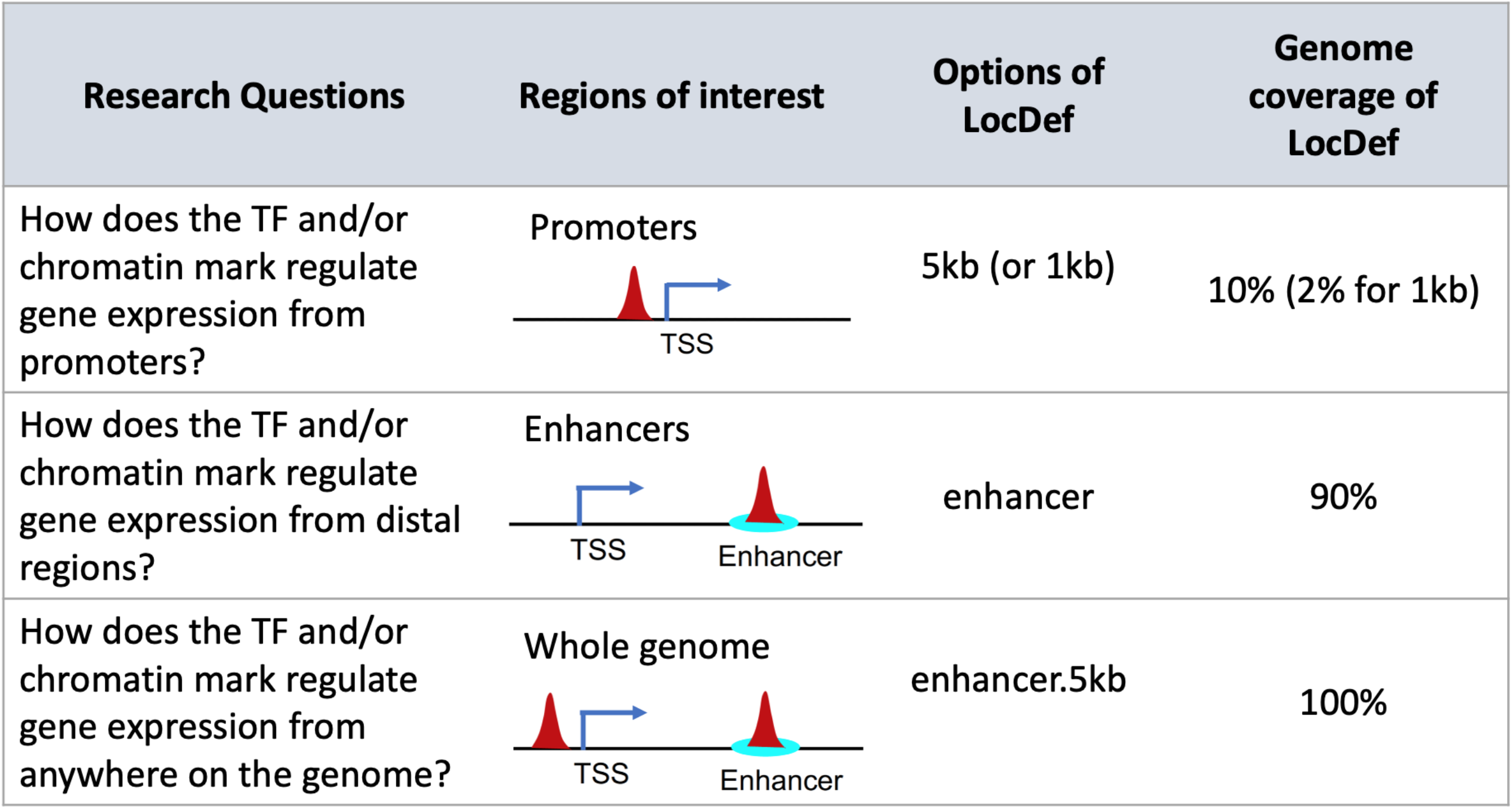
User guidelines for selecting an appropriate enhancer-to-gene assignment method (LocDef) for GSE testing. Depending on the specific research questions, three types of LocDefs can be selected for GSE testing from the *chipenrich* R package: 1) “5kb” or “1kb” for promoter regulation, 2) “enhancer” for distal regulation, and 3) “enhancer.5kb” for whole genome regulation. Different LocDefs have different genome coverages as shown in the last column. Options in other GSE testing software for genomic regions will differ. We no longer recommend using nearest TSS method for Poly-Enrich analysis.

## DISCUSSION

A greater appreciation of the central role that distal regulatory elements play in genetic diseases and cancers has motivated a multitude of enhancer studies. As a result of the increasing availability of functional genomics data, growing attention has been paid to matching Enhancer-Target Gene pairs (ETG) in the field of computational biology and genomics. Over the past decade, a variety of algorithms and tools have been developed by leveraging multiple genomic features and functional data, as recently reviewed in [51]. Briefly, they can be categorized into four groups: *1)* correlation-based (e.g. Thurman *et al* [38], PreSTIGE [52], ELMER [53, 54], etc.); *2)* supervised learning-based (e.g. IM-PET [55], TargetFinder[56], McEnhancer[57], etc.); *3)* regression-based (e.g. RIPPLE[58], JEME[59], FOCS[60], etc.); and *4)* score-based methods (e.g. EpiTensor[61], GeneHancer[62] and PEGASUS [63, 64]). Although these algorithms have significantly advanced our knowledge of ETGs, they are affected by one or more of the following issues: *1)* the lack of a genome-wide exhaustive reference list of enhancers; *2)* the lack of a large gold standard which is required for supervised learning algorithms, i.e. experimentally-validated true positive and true negative enhancer-target gene pairs, and *3)* the lack of a systematic evaluation of their reliability and generalization in various cell types. To overcome these issues, we developed a gold standard-free approach to generate and prioritize comprehensive sets of ETGs based on their performance in the interpretation of regulome data.

In this study, we identified a best set of **En**hancer to **T**arget gene **Def**initions (**EnTDefs**) by investigating and evaluating all possible combinations of existing reliable sources for human enhancer location definitions and enhancer-target gene pair definitions across various cell types. Purposely, we coupled EnTDefs with GSE testing to systematically evaluate their performance when interpreting regulome data. By carefully selecting datasets of high quality and resolution, we explored ENCODE ChromHMM, DNase-seq, FANTOM5 and Thurman datasets for enhancer regions and ENCODE ChIA-PET interactions, Thurman DHS correlation-based, FANTOM5 and ENCODE ChIA-PET CTCF loops-based enhancer-target gene interactions. We also systematically evaluated the performance of all possible combinations of datasets when applied on ENCODE TF ChIP-seq data in GO GSE testing and compared the enriched GO terms with the curated TF GO annotations (TF-annotated GO BP terms by the GO database). In contrast to the statistical modelbased or machine learning-based algorithms as described above, our approach integrates various data sources and directly couples the EnTDefs with GSE testing for a systematic evaluation, resulting in an EnTDef with maximally balanced sensitivity and specificity (assessed by F1-score). Our approach to generating EnTDefs is assumption-free and independent of true positive/negative pairs, but based on a systematic evaluation using GSE testing. The results demonstrate that the DNase-seq and FANTOM5 enhancers with the integrated enhancer-target gene pairs from ChIA-PET, Thurman and FANTOM5 interactions performed best, suggesting that both chromosome accessibility and conformation, as well as transcriptional correlation, are beneficial for identifying enhancer-target regulatory relationships.

The nearest distance method, which naively assigns genomic regions of interest to the nearest gene, is commonly used in GSE for regulome data. Our analysis showed that this naïve approach commonly fails to identify certain functions, such as those related to chromosome organization, which lead to false negatives, whereas our top EnTDefs can successfully identify these long-range enhancer regulatory functions. These findings re-confirmed that universally assigning cis-regulatory elements to the gene with the nearest TSS is problematic, resulting in misleading and/or incomplete functional interpretation.

Our EnTDefs were generated by leveraging different genomic data across >500 cell types and can be applied to different cell types, demonstrating performance comparable to their cell-type-specific counterparts. Our top integrated EnTDef based on many cell types represents a comprehensive set of enhancer regions (only a subset of which will be active in any one cell type); our data indicate this performs well because current cell-type-specific enhancer-target genes (ETGs) are not yet sufficiently comprehensive (except for a few cell types such as GM12878). Research performed on cancer samples, less commonly used cell lines, and other complex tissue samples will greatly benefit from this integrated EnTDef. While cell-type-specific ETGs are important for studying regulation at specific locations, our results demonstrate that for genomewide approaches such as GSE, the comprehensiveness outweighs the need for specificity.

Besides DNase-seq, ChIA-PET, CAGE-seq, and RNA-seq data, Hi-C and eQTL data are also used to infer ETG [62, 65]. However, we found that current Hi-C data often have insufficient resolution, with genomic windows being a few to several kb wide due to low coverage, and high quality Hi-C data is not available for nearly as many cell types as the other approaches. Although eQTL data is available for many tissues and cell types, it is similarly restricted by limited population diversity and low resolution. The tissue-specific eQTL data from the GTEx project [66] is widely used, however, it was generated for only 49 tissues from <1000 donors with the majority being Caucasian (84.6%), making it difficult to apply to other tissues/populations. In addition, eQTL data is highly correlated with the linkage disequilibrium, and thus its resolution is associated with the size of haplotype blocks, which is highly variable across populations (on average ~10kb) [67], whereas enhancers are usually short genomic regions (50-1,500bp). Due to this low resolution of Hi-C and eQTL data, we excluded them from our analysis. On the other hand, another data-integration method, HACER, was recently developed [68], which utilized the nascent eRNA information from GRO-seq and PRO-seq, along with the FANTOM5 CAGE-seq data, to identify cell-type-specific ETGs. In future work, we will incorporate the GRO-seq and PRO-seq data deposited in HACER and evaluate if the new datasets can further boost our EnTDef performance when coupled with GSE.

In conclusion, we identified a best set of enhancer-target gene pairs (EnTDef) by leveraging existing data sources of chromosome accessibility and/or conformation and transcriptome data across numerous cell types, which significantly improved the biological interpretation of distal regulation in GSE compared to assigning genomic regions to the nearest gene. Our approach performs well across a wide range of cell types, making it feasible to apply on extensive genomic data sets. The limitations of our EnTDef are inherited from the existing data sources, including low genome coverage, low resolution, and small number of cell types with good quality ChIA-PET data. With the continued growth in volume of functional genomics data and advances in data quality and resolution, we expect further improvement of our EnTDef in the future.

### Conclusions

In summary, we provide an optimized enhancer-to-target gene assignment approach, which is critical for interpreting genome-wide regulatory data. This study has important implications for which type of enhancer-target gene methods are most accurate, and the relative importance of comprehensiveness versus cell-type specific accuracy. To the best of our knowledge, there is currently no such a comprehensive resource of distal regulatory region-to-target gene links which are feasible to apply on various types of regulome data (eg.ChIP-seq, ATAC-seq) regardless of cell types.

## METHODS

### Generation of general enhancer-target gene definitions

We generated genome-wide definitions of human distal enhancer locations and their target gene assignments for the hg19 genome using all possible combinations of the below enhancer location methods and enhancer-gene linking data (**Figure 1A**: Enhancer, Extension, Enhancer-target gene link and Additional links). These are based on enhancers from: *1)* “ChromHMM”: ENCODE ChromHMM UCSC tracks (9 cell types) [43], *2)* “DNase-seq”: DNase hypersensitive sites (DHSs) from 125 cell types processed by ENCODE [38], *3)* “FANTOM5”: Cap Analysis Gene Expression (CAGE) experiment-derived enhancers across 421 distinct cell lines/tissue/primary cells from FANTOM5 project [37, 44, 45], and/or *4)* “Thurman”: distal and non-promoter DHS within 500 kb of the correlated promoter DHSs from 79 cell types, referred to as the first author of the publication [38]. Since our motivation was to identify the target genes of distal regulatory elements that do not have clear target genes based on close proximity to a TSS, we constrained the enhancer regions to be outside of 5 kb from a transcription factor start site (TSS) by trimming the bases from the above defined enhancers overlapping with the 5kb windows of TSSs. The hg19 TSS locations were obtained from the Bioconductor *chipenrich* package version 3.5.0 [42]. To identify target genes, we used: *1)* “ChIA” method: enhancer and gene interactions identified by ChIA-PET2 using default parameters [69] from 10 ChIA-PET datasets of 5 cell types (**Supplementary Table S3**) [46, 47], *2)* “Thurman” method: the enhancer and promoter interactions identified by Thurman *et al,* which were defined by high correlation (r > 0.7) between cross-cell-type DNase I signal at each DHS position and all promoters within ±500 kb [38], *3)* “FANTOM5” method: the regulatory targets of enhancers predicted by correlation tests using the expression profiles of all enhancer-promoter pairs within 500kb[45], and *4)* “Loop” method: any possible interactions between enhancers and genes that are encompassed within in a RAD21, cohesin and/or CTCF ChIA-PET loop with convergent CTCF motifs [48], and depending on the number of genes included in the loop, this method was referred to as “L1” (one gene), “L2” (≤ two gene) or “L3” (≤ three genes) (**Figure 1B**).

All possible combinations of the above, allowing multiple at a time, defined 465 of the **En**hancer to **T**arget gene **Def**initions (EnTDefs) (**Figure 1A, B)**. In addition, to increase the genome coverage, we tested extending the enhancer regions to 1kb (i.e. “enhancer extension”, 500 bp extension at both sides of the midpoint), and assigning regions outside of enhancers and promoters (within 5kb of a TSS) to the gene with the nearest TSS (i.e. “nearest_all” additional links). The additional combinations using these options brought us to a total of 1,860 distinct EnTDefs.

### Evaluation of enhancer-target gene definitions

To evaluate the performance of each individual EnTDef, we performed Gene Ontology (GO Biological Processes [GOBP]) enrichment testing using Poly-Enrich [70] in the *chipenrich* Bioconductor package[42] on 87 ChIP-seq datasets of 34 TFs selected from the tier 1 ENCODE cell lines (**Supplementary Table S4**). We then compared the significantly enriched GOBP terms with the GO BP annotations of each TF (i.e. the GOBP terms assigned to the 34 TFs by the GO database, excluding the terms with <15 or >2000 assigned genes) (**Figure 1C**: Evaluation of the Enhancer-Target gene Definition), to identify the EnTDefs with greatest concordance. The assumption of this approach, used previously in[70, 71], is that TFs tend to the regulate genes in the biological processes to which they belong, and thus greater overlap with TF GO BP annotation indicates more accurate enrichment results, and thus more accurate peak-to-gene assignments. For a full justification of this, see Supplementary Methods. To minimize runtime for the initial pass analysis, we used the PE.Approx method (an approximate version of Poly-Enrich[70], see Supplementary Methods and **Supplementary Figure S7**). To alleviate the bias caused by the unbalanced number of positive and negative assignments, we generated the same number of true negative assignments for each TF as there were positive by randomly selecting GOBP terms from the set that were not assigned to the particular TF, and excluding the offspring terms and their siblings of the assigned terms (hereafter called “true negative” terms, depicted in **Supplementary Figure S8A**). In order to control for the confounding of GOBP size (i.e. the number of assigned genes to each GOBP term), random sampling was performed among the negative terms of comparable size to the corresponding true positive term (bin size = 20). In each sampling, the PE results were assessed by the number of true positive (TP), false positive (FP), true negative (TN) and false negative (FN) GOBP terms according to the following definitions: *1)* TP: the number of GOBP terms that were significantly enriched (FDR < 0.05) and assigned to the TF by the GO database; *2)* FP: the number of GOBP terms that were significantly enriched (FDR < 0.05), but not assigned to the TF by the GO database; 3) TN: the number of GOBP terms that were not significantly enriched (FDR > 0.05 or “depleted”) and also not assigned to the TF; and 4) FN: the number of GOBP terms that were not significantly enriched (FDR > 0.05, or “depleted”), but assigned to the TF. The *F1 score* 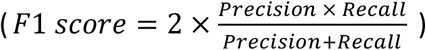 was calculated to measure the overall performance of an EnTDef for a TF. We repeated the sampling process 10 times, and took the average *F1 score* for each EnTDef and TF. The average of these F1 scores across TFs provided the final ranking for each EnTDef.

To assess the robustness of our approach, we also evaluated the performance of EnTDefs using more conservative GO annotations, in which GOBP assignments based on “automatically assigned, inferred from Electronic Annotation” (IEA) were excluded, thus minimizing false annotations in GOBP. For the positive GO annotations we used only the leaf GO terms (the lowest level in the GO hierarchical tree) and their parent and grandparent GO terms, while the negative terms were sampled from all other terms, excluding positive terms, and ancestors of positive terms, siblings of ancestors of positives, as well as offspring of positives (depicted in **Supplementary Figure S8B**).

After ranking all EnTDefs in descending order by their average F1 scores (**Figure 1C**: Rank of EnTDef), we identified the set of best EnTDefs. Paired Wilcox signed-rank tests were performed to compare the F1 score of the 1^st^ ranked EnTDef with each of the sequential ones, and the rank at which the EnTDef showed significantly lower F1 score than the 1^st^ ranked one (p < 0.01) was selected as the cutoff. The EnTDefs ranking above the cutoff were defined as the best set of EnTDefs. In addition, we performed the same F1 score evaluation of previously-defined methods for genomic region-to-gene assignments, termed gene **Loc**us **Def**initions (LocDefs, see **Supplementary Figure S1** for details) that do not use “smart” enhancer-target links (i.e., “>5kb”: distal regions assigned to the gene with the nearest TSS; “<5kb”: regions within 5kb of a TSS assigned to the gene with that TSS; and “nearest TSS”: all regions assigned to the gene with the nearest TSS). These LocDefs are used by Poly-Enrich in the *chipenrich* R Bioconductor package [42], and represent the current standard practice for enhancer-to-gene assignments for gene set analysis. The F1 scores were compared between each EnTDef and the distal nearest distance (“>5kb”) LocDef by Wilcoxon signed-rank tests. We also evaluated and compared two commonly used Gene Set Enrichment (GSE) testing methods, Fisher’s Exact Test (FET) and GREAT [39], which were implemented by the R *chipenrich* package using the FET and binomial method respectively, coupled with the “5kb” LocDef. To obtain a final assessment, a second round of GSE testing using Poly-Enrich (*PE.Exact* method; see Supplementary methods for details) was applied on the subset of EnTDefs which significantly outperformed the nearest distance assignments (>5kb LocDef), and the average F1 scores were calculated and used to refine the final ranking of EnTDefs.

### Validation of the EnTDefs with ChIP-seq data from different cell types

To further evaluate the performance of EnTDefs in different cell lines, we selected 13 additional ENCODE ChIP-seq datasets from four non-tier 1 ENCODE cell lines (A549, HEPG2, HUVEC and NB4), which contain ChIP-seq experiments for at least three TFs in each cell line (**Supplementary Table S5).** In comparison to the 87 *evaluation* ChIP-seq peak sets from ENCODE tier 1 cell lines (GM12878, H1HESC and K562), these 13 datasets are the *test* datasets. The six TFs (*C-JUN, C-MYC, CEBPB, CTCF, MAX* and *NRSF*) assayed by the 13 *test* datasets were also included in the 87 *evaluation* datasets. The top 10 best EnTDefs were evaluated using the *PE.Exact* method as described above and these 13 test ChIP-seq datasets. For each EnTDef, the average F1 score across the 13 ChIP-seq datasets was calculated and compared with the average F1 score generated using the evaluation ChIP-seq datasets (n=16) of the corresponding TFs.

### Generation of cell-type-specific EnTDefs

We used “ChIA” and/or “L”-derived enhancer-to-gene assignment methods (**Figure 1B**) to generate cell-type-specific enhancer-target gene definitions, hereafter called CT-EnTdefs. Since the enhancer-gene linking data defined by Thurman and FANTOM5 datasets were non-cell-type specific, we did not include these. The cell types were selected based on the availability and quality of cell-type specific ChIP-seq and ChIA-PET data in ENCODE. As shown in **Supplementary Table S5**, four cell types were selected: GM12878 (tier 1), H1-hESC (tier 1), K562 (tier 1) and MCF7 (tier 2). The multiple ChIA-PET datasets were combined for each cell type. All combinations of enhancer location definitions, along with ChIA, L1 (or L2 or L3) enhancer-gene assignment methods, with or without enhancer location extension and “nearest_all” addition, were used to generate the CT-EnTDefs, resulting in a total of 630 CT-EnTDefs for each of the 4 cell types.

### Evaluation of CT-EnTDefs

To evaluate the performance of the CT-EnTDefs, we performed GSE testing of Gene Ontology (GO Biological Processes [BOBP]) using Poly-Enrich [70] on the TF ChIP-seq peak sets of the same cell type from which each CT-EnTDef was generated (**Supplementary Table S5**. See details as described above). For comparison, we also applied GSE testing on the same TF ChIP-seq peak sets using the corresponding general EnTDefs (i.e. not cell-type specific, using the same enhancer regions and target gene link methods as those of the comparative CT-EnTDef), as well as the CT-EnTDef from a different cell type (Figure 3C, i.e. MCF7 CT-EnTDefs were applied on GM12878 TF ChIP-seq peaks, GM12878 CT-EnTDefs on H1hESC peaks, H1hESC EnTDefs on K562 peaks, K562 EnTDefs on MCF7 peaks). For each TF, the average *F1* scores across all evaluated EnTDefs were calculated and compared between using the respective CT-EnTDef, general EnTDef and different cell type CT-EnTDef. For each cell type, Pearson’s correlation test was used to evaluate the pairwise correlation among the F1 scores, and Wilcoxon Rank-Sum test was used to compare their differences. Finally, the overall performance of CT-EnTDefs, general EnTDefs and different cell type CT-EnTDefs were assessed using the average F1 scores across all evaluated EnTDefs and TFs in each cell type.

### Testing for functions that have significantly more or fewer interceding genes between enhancers and their target genes

We investigated the number of interceding genes between an enhancer and its target gene(s) (i.e. genes between the entire region of an enhancer and the target gene in an EnTDef, depicted in **supplementary Figure S6**), and ranked all target genes based on their average number of interceding genes. By definition of nearest distance enhancer-target gene assignment (e.g. >5kb LocDef), the bottom genes with low numbers of interceding genes are most likely to be correctly assigned to their enhancers, while the top ranked genes with high numbers of interceding genes are least likely assigned to their true enhancers. We used the best performing EnTDef without “nearest_all” addition, defined by DNase-seq plus FANTOM5 enhancers and ChIA, Thurman and FANTOM5 enhancer-target gene link methods, as an example for this analysis. Gene Ontology (GO) enrichment testing was performed by *LRpath[50]* using GO Cellular Component (CC), Biological Process (GOBP), and Molecular Function (MF) terms of size ranging from 10 to 1000 genes. The rank-based inverse normal transformation (INT) implemented by the *rankNorm* function in R package *RNOmni*[72] was applied to the average number of interceding genes to have approximately normally distributed scores. *LRpath* took the genes that were linked to at least one enhancer and their exponential transformed INT scores as the input (the input scores were log transformed internally by *LRpath* program) and performed logistic regression-based enrichment testing on each GO term. The significant GO terms (FDR < 0.05) with positive coefficients indicate the functions enriched in genes with less interceding genes (lower ranked), while those with negative coefficients are functions enriched in genes with more interceding genes (higher ranked). For reporting purposes, we filtered out closely related GO terms, using the *GO.db* R package [73] to determine relationships among significant terms. A GO term was filtered if one or more of its parents, children or siblings had a higher rank in the list [74].

To determine the robustness of the results, we performed the same analysis for all top 10 best performing EnTDefs without “nearest_all” addition. The enrichment results were combined across EnTDefs for each GO term by taking the Harmonic Mean (HM) p-values[75]. The significant terms were extracted using FDR-adjusted HM p-values (HM FDR < 0.05), followed by redundant term filtering as described above.

### Identification of common false positive and false negative enrichment results by nearest distance enhancer-target gene assignment

Next, we examined the false positive (FP) and false negative (FN) GO terms identified using the nearest distance assignment (>5kb LocDef), that were correctly identified using the ‘smart’ enhancer-target gene assignments. To do this, we compared the Poly-Enrich [70] GSE results of the 87 evaluation ChIP-seq datasets using >5kb LocDef for nearest distance to those of the best performing EnTDef (defined by DNase-seq plus FANTOM5 enhancers and ChIA, Thurman and FANTOM5 enhancer-target gene link methods with “nearest_All” addition) using the known TF-GO BP annotated terms (i.e. GO annotation). The FP and FN GO BP terms were defined according to the following criteria: *i)* FPs: GO terms identified as significantly enriched (FDR < 0.05) by the >5kb LocDef, but not significant (FDR ≥ 0.05) by EnTdef and not annotated by the GO database; *ii)* FNs: GO terms identified as insignificant (FDR ≥ 0.05) by the >5kb LocDef, but significantly enriched (FDR < 0.05) by EnTDef and also annotated by the GO database. GO terms were removed if they were annotated to < 4 (<10%) or > 30 (>90%) of the 34 used TFs to ensure the possibility of having both true and false annotations present among the TFs, and only terms with < 1000 total annotated genes were used in this analysis. For each GO term, the number of FPs and FNs among the 34 TFs, detected using ‘>5kb’, were summarized, and the proportion of FP and FN were calculated using the total number of positive TFs (FDR < 0.05) or negative TFs (FDR ≥0.05) by >5kb LocDef as the denominator, respectively. To summarize the significance level of each FP or FN term in GSE testing, a meta-p-value was generated using Fisher’s method [76] across the annotated TFs using the raw p-values from the >5kb LocDef or EnTDef approach, respectively. Finally, the FP (or FN) GO terms were ranked by the proportion of FP (or FN) decreasingly and meta-p value increasingly, followed by redundant GO term filtering as described above.

## DATA AVAILABILITY

The top performing EnTDef and EnTDef.plus5kb have been included in the Bioconductor package *chipenrich* (42) and the ChIP-Enrich website (https://chip-enrich.med.umich.edu). The peak-to-gene assignment functionality provided by our GSE Suite (http://gsesuite.dcmb.med.umich.edu) allows users to select all possible combinations of enhancer and/or enhancer-to-gene link methods (as described in this study) and obtain the gene assignments for user uploaded genomic regions based on the selected sources and methods. Genomic regions can also be assigned to target genes based on other approaches (e.g. promoters, exons or anywhere in the genome using the nearest distance method).

## ACKNOWLEDGEMENTS

T.Q. performed the bioinformatics analysis and wrote the manuscript. C.L. contributed the GSE testing. R.C. and P.O. contributed the data collection and EnTDef generation. H.Y., H.Z. and S.W. contributed specific bioinformatics analysis. S.P. developed the GSE Suite. A.P.B. contributed to manuscript writing and provided analysis suggestions. M.A.S. conceived, supervised the study, participated in data interpretation and manuscript writing.

## Supplementary figures

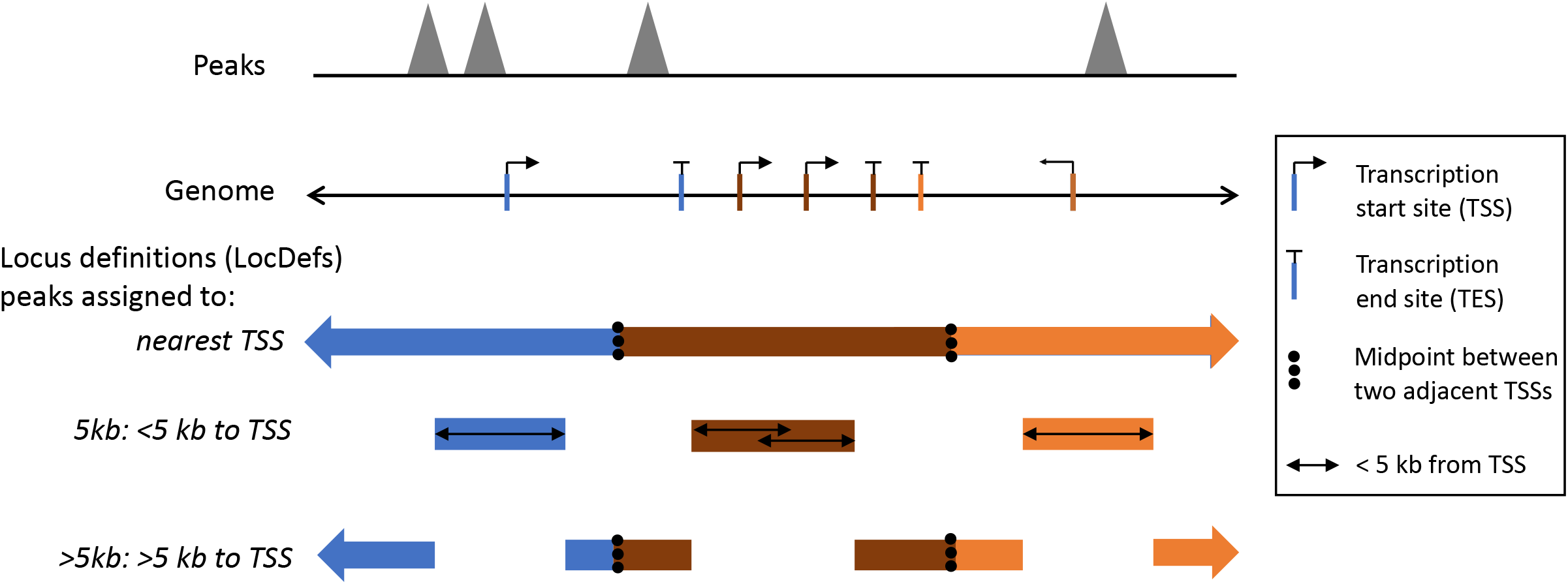

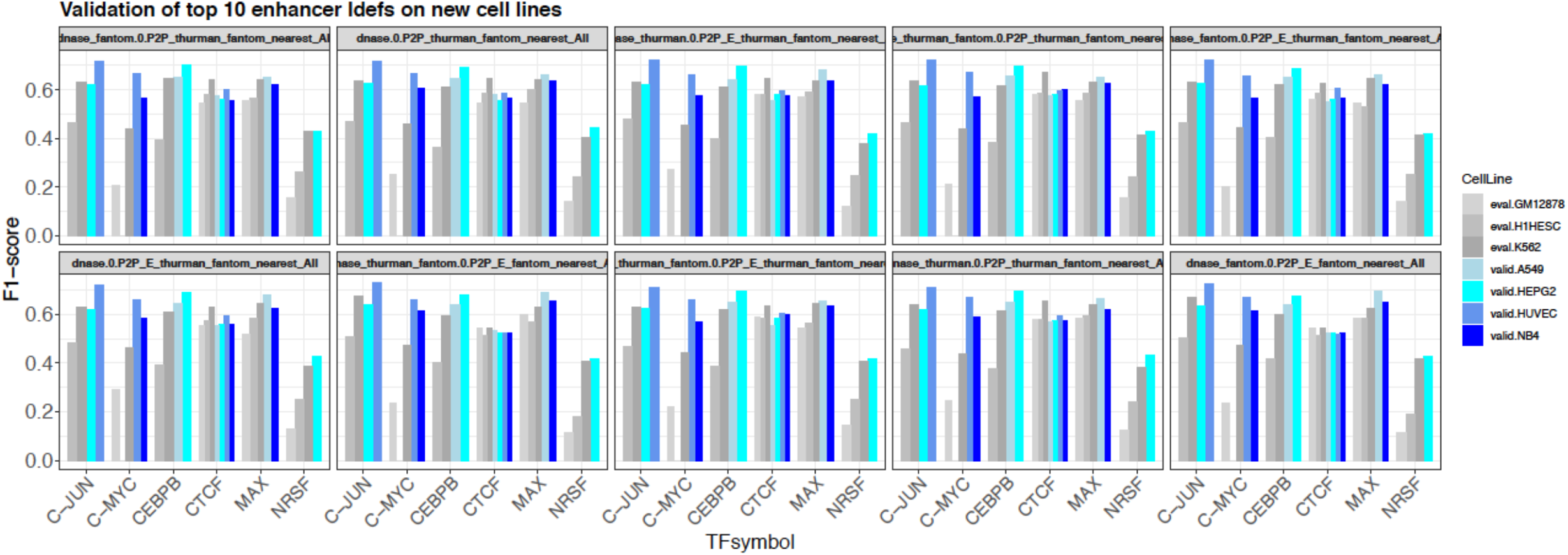

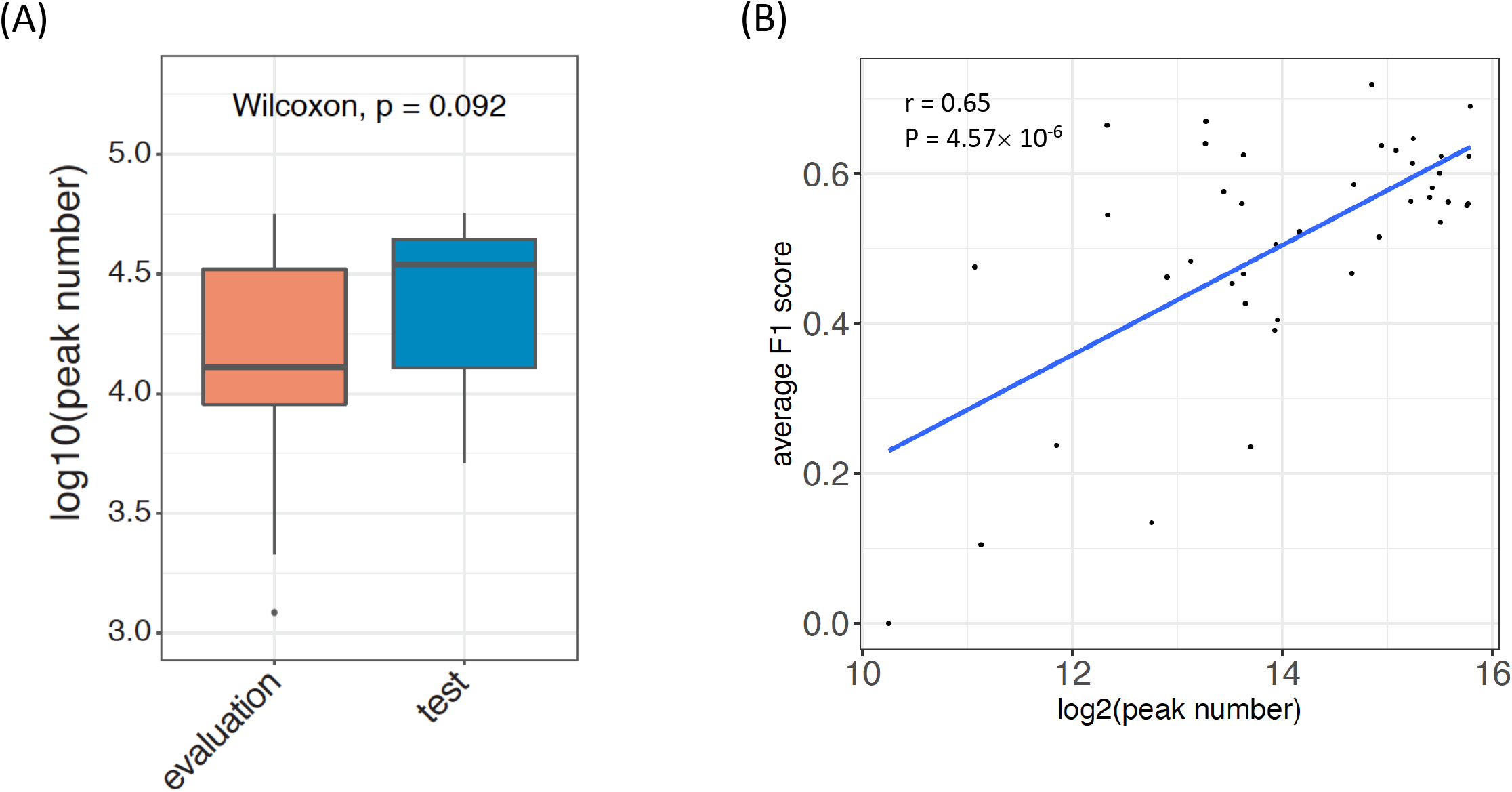

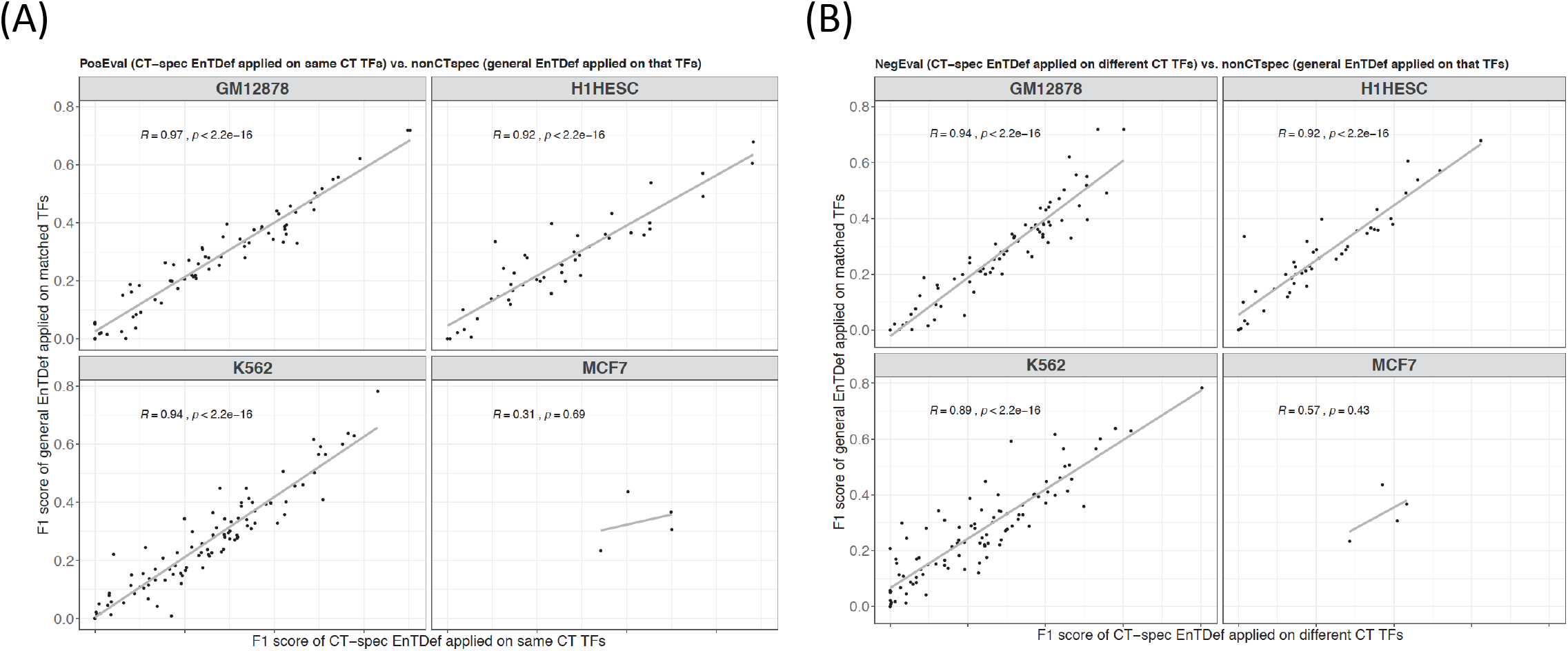

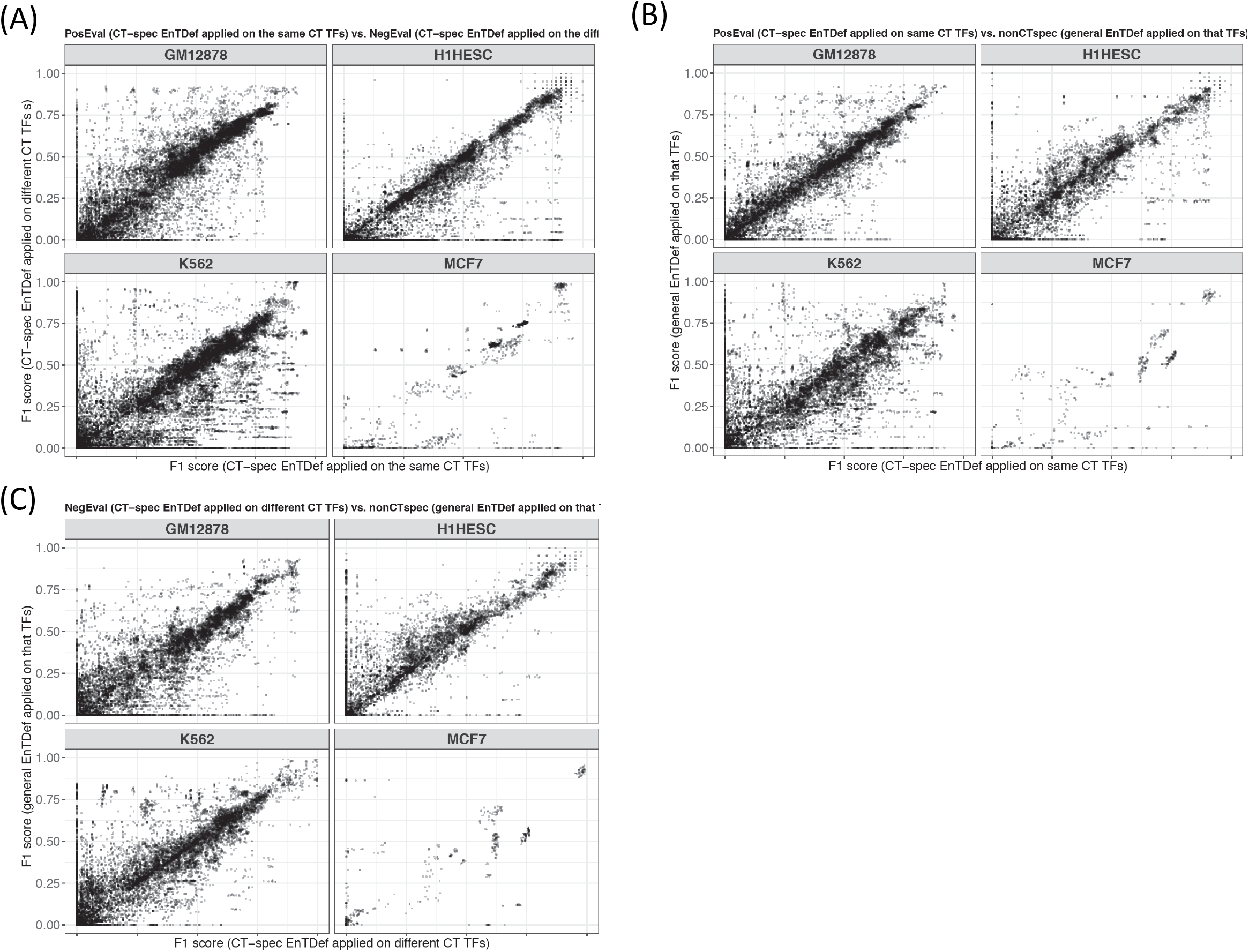

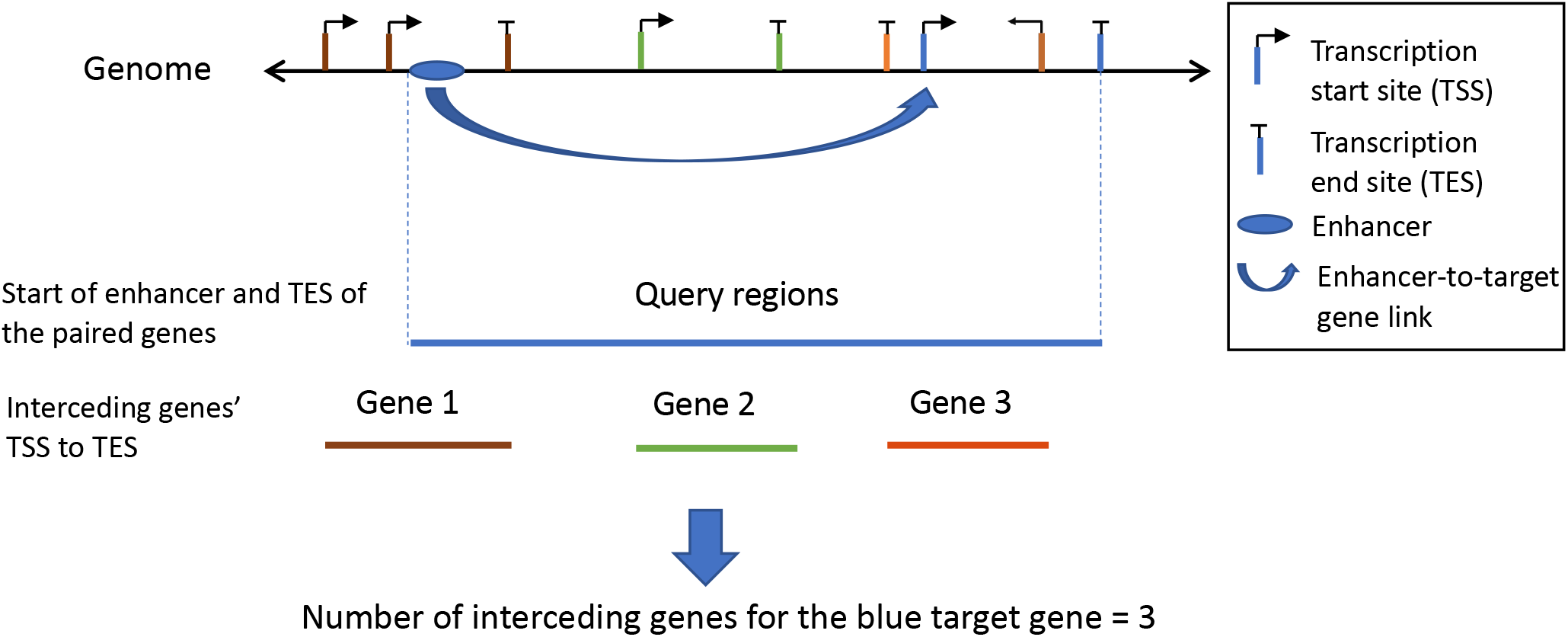

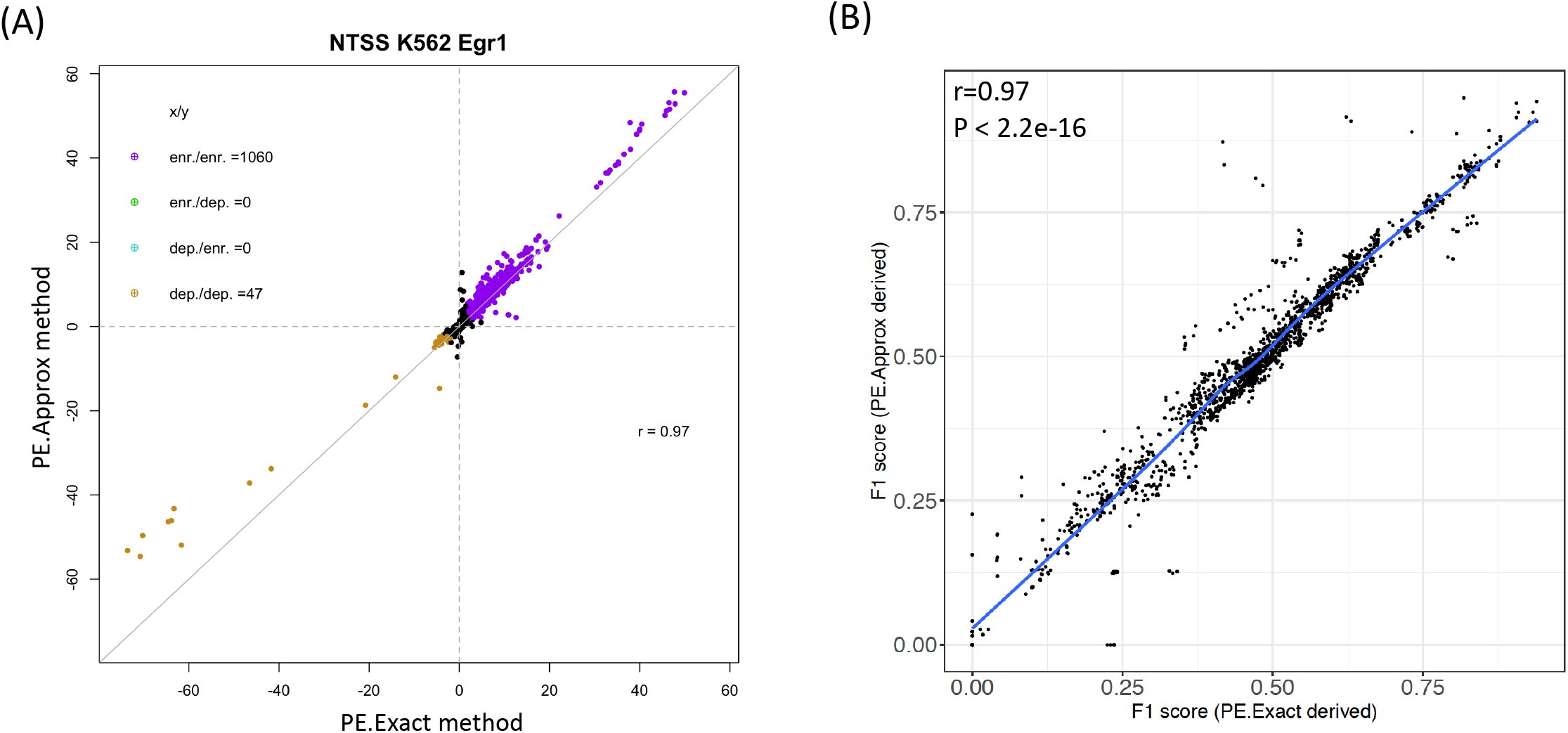

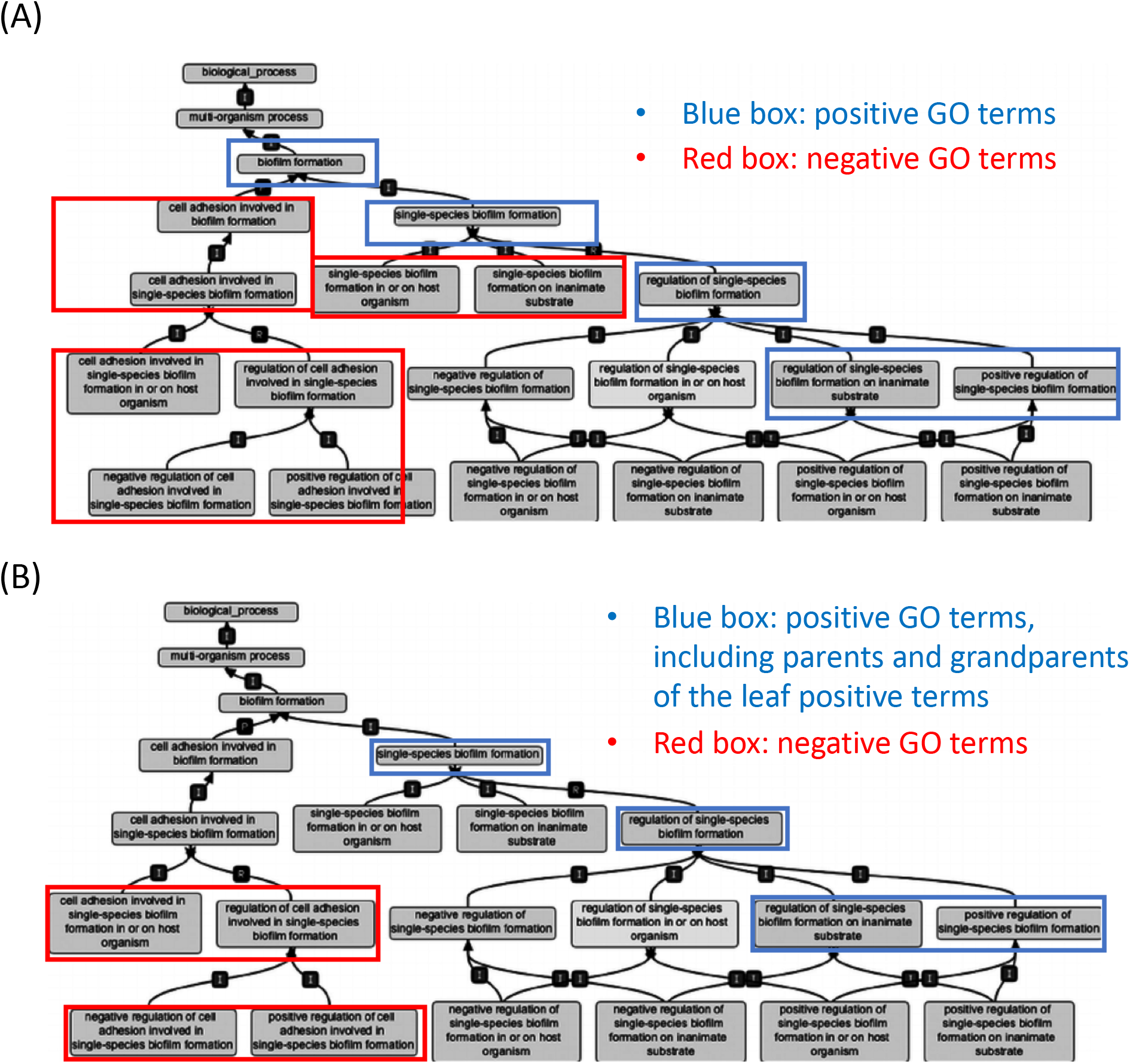

