## Supplementary methods for "Comprehensive enhancer-target gene assignments improve gene set level interpretation of genome-wide regulatory data"

### *Comparison between Poly-Enrich approximate (PE.Approx) and exact (PE.Exact) methods in GSE testing*

For the first round of evaluating **En**hancer to **T**arget gene **Def**iniations (EnTDefs), we applied an faster, approximate version of Poly-Enrich (PE) test, which utilized the score test instead of likelihood ratio test. Compared to the Poly-Enirch likelihood ratio test (i.e. PE.Exact method), the Poly-Enrich score test (i.e. PE.Approx method) requires the least amount of computation time in exchange for lacking power in negative binomial families [1]. However, the score test is a good approximate test for situations when one needs a large amount of preliminary results. We used the *glm.scoretest* function from the *statmod*[3] package to compute the score test for PE.Approx method. The PE.Approx runs over 30 times faster and is reasonably concordant with the PE.Exact for enriched GO terms, but less similar for depleted GO terms (**Supplementary Figure S7A**). The *F1 score* of the top 19 best EnTDefs derived from the PE.Approx GSE method was highly correlated to that derived from the PE.Exact method (Spearson's correlation $r = 0.97$, $p < 0.00001$, **Supplementary Figure S7B**).

### *Using GO-to-TF assignments by the GO database as true positives to evaluate the enhancer-target gene definitions in GSE testing*

To evaluate the performance of the enhancer-target gene definitions (EnTDefs), we performed Poly-Enrich[4] GSE testing on the evaluation ChIP-seq datasets (87 ENCODE ChIP-seq of 34 TFs) using EnTDef-defined genomic region-to-gene assignments, and compared the significantly enriched gene sets for each transcription factor (TF) with the ones the TF is likely to regulate. Since no gold standard is available for this, we used the Gene Ontology Biological Process (GO BP) terms that each of the 34 TFs were assigned to by the GO database [5, 6], and extracted the GOBP-to-gene assignments from the human annotation Bioconductor package org.Hs.eg.db [7]. We reasoned that TFs tend to regulate genes in the biological processes to which they are assigned in GO [8-11]. Since the GO BP terms describe "the larger processes, or 'biological programs' accomplished by multiple molecular activities"[6] and the function of a TF is to regulate genes that coordinate a common biological process, it is logical to assume that TFs tend to regulate genes in the GO BP to which they belong. Although a TF may not regulate all of their assigned GO terms in every cell type, we assume that higher concordance with this set (in terms of sensitivity and specificity) corresponds to superior results. To alleviate the bias caused by the unbalanced positive and negative assignments (i.e. each TF only regulates a very small percent of the total GO terms), we generated the same number of the true negative assignments for each TF by randomly selecting the GOBP terms (see Methods).

**Supplementary table titles**

**Table S1.** The F1 score paired-testing results between the 1$^{st}$ ranked EnTDef vs. the sequential ranked ones using paired t-tests and Wilcoxon signed-rank tests.

**Table S2.** The nine ChIA-PET datasets used for generating cell-type-specific EnTDefs (CT-EnTDefs) and number of TFs assayed by ENCODE ChIP-seq in each particular cell type, which were used to evaluate the performance of the CT-EnTDefs.

**Table S3.** ChIA-PET datasets used by “ChIA” and “Loop” methods to assign enhancer to target genes in a cell-type independent manner (general EnTDefs).

**Table S4.** The 87 ENCODE ChIP-seq datasets used for EnTDef evaluation (evaluation ChIP-seq).

**Table S5.** The 13 ENCODE ChIP-seq datasets from 4 different cell lines (testing ChIP-seq).

**Supplementary figure legends**

**Supplementary Figure S1. Illustration of different types of Locus Definitions (LocDefs) used in this study.** ChIP-seq peaks (top) are assigned to genes if they are located within a chosen LocDef, including: “nearest TSS”, “<5kb to TSS” and “>5kb to TSS”.

**Supplementary Figure S2. Bar plots of *F1 scores* for each cell type and TF among the evaluation and testing ChIP-seq data sets.** Each panel represents one of the top 10 EnTDefs. Cell types in the evaluation dataset are greyish, while those in the testing dataset are bluish.

**Supplementary Figure S3. Number of peaks in the ChIP-seq datasets and their correlation with F1 score.** (A) Boxplots of the number of peaks in the evaluation and testing ChIP-seq data. (B) The correlation between the number of peaks in evaluation/testing ChIP-seq datasets (log2 scale) and the average *F1 score* across the top 10 best EnTDefs.

**Supplementary Figure S4. Correlation of average *F1 scores* for a TF across EnTDefs.** (A) The correlation between average F1-scores calculated on a TF in a particular cell type using CT-EnTDefs of the matched cell type (x-axis) and the ones calculated on the same TF using general EnTDefs (y-axis). (B) The correlation between average F1-scores calculated on a TF in a particular cell type using CT-EnTDefs of a different cell type (x-axis) and the ones calculated on the same TF using general EnTDefs (y-axis). Each dot represents an average F1-score of a TF across EnTDefs, and each panel is one of four cell types (GM12878, H1HESC, K562 and MCF7) for which the CT-EnTDefs were created and evaluated respectively.

**Supplementary Figure S5. Correlation of *F1 scores* for each TF and EnTDef pair.** (A) The correlation between F1-scores calculated on a TF in a particular cell type using a CT-EnTDef of the matched cell type (x-axis) and the ones calculated on the same TF using a CT-EnTDef of a different cell type (y-axis). (B) The correlation between F1-scores calculated on a TF in a particular cell type using a CT-EnTDef of the matched cell type (x-axis) and the ones calculated on the same TF using a general EnTDef (y-axis). (C) The correlation between average F1-scores calculated on a TF in a particular cell type using a CT-EnTDefs of a different cell type (x-axis) and the ones calculated on the same TF using the general EnTDef (y-axis). Each dot represents a F1-score of a TF and EnTDef pair, and each panel is one of four cell types (GM12878, H1HESC, K562 and MCF7) for which the CT-EnTDefs were created and evaluated respectively.

**Supplementary Figure S6. Illustration of interceding gene definition.** The interceding genes were defined as the genes with any part of the gene body falling in the query region (i.e. the genomic regions between the farthest positions of an enhancer and its target gene pair).

**Supplementary Figure S7: Comparison between Poly-Enrich approximate and exact methods**

(A) The representative GSE result of transcription factor (TF) *EGR1* in the K562 cell line: all GO terms are generally concordant with the score test (PE.Approx method) being slightly more conservative, but the depleted GO terms tend to deviate more from the PE.Exact method. (B) The correlation of *F1 score* of the top 19 best EnTDefs between the GSE result derived from the PE.Approx method and that derived from the PE.Exact method. Each dot represents a GSE result of a particular TF using one of the 19 EnTDefs.

**Supplementary Figure S8. Illustration of positive and negative GOBP terms assigned to TFs by GO with the number of annotated genes ≥ 15 and ≤ 2,000.** (A) Positive terms include the lowest level of assigned GOBP (leaf terms) and all of their ancestors; negative terms include the terms outside of positive ones, excluding the leaf terms and their siblings and offspring of the assigned terms. (B) More conservative sets of positive and negative terms: positive terms include assigned leaf terms and their parent and grandparent terms, excluding the ones assigned by IEA (automatically assigned, inferred from Electronic Annotation); negative terms include the terms outside of the positive ones, excluding the ancestors of positives, siblings of positives' ancestors, and offspring of positives.
